# Functionally Relevant and Reliable Brain Stimulation Targets for Enhancement of Novel Word-Learning

**DOI:** 10.1101/2025.11.04.686295

**Authors:** Harun Kocataş, Mohamed Abdelmotaleb, Leonardo M. Caisachana Guevara, Filip Niemann, Alireza Shahbabaie, Robert Malinowski, Steffen Riemann, Dayana Hayek, Daria Antonenko, Antoni Rodriguez-Fornells, Agnes Flöel, Marcus Meinzer

## Abstract

Linking word-forms and their meanings is central to language learning. Transcranial direct current stimulation (tDCS), has shown potential to enhance this process, but with variable effects. This study aimed to (1) identify reliable and functionally relevant tDCS target brain regions to enhance novel-word learning and (2) assess test–retest reliability (TRR) of behavioral and imaging outcomes. Twenty healthy individuals completed two functional magnetic resonance imaging (fMRI) sessions using parallel task versions. Participants learned picture-pseudoword associations across six learning blocks. Behavioral learning was analyzed using linear-mixed-models. Whole-brain and region-of-interest (ROI) analyses examined learning-related activity changes and their behavioral relevance. TRR was assessed using intraclass correlation coefficients (ICCs). Participants successfully acquired the novel-word forms, indexed by increased accuracy and faster latency across stages. Behavioral outcomes showed good-to-excellent TRR. The task elicited robust language-learning related activity and activity changes across stages were correlated with learning success. Task-related activity was variable, but voxels within significant clusters (∼81%) and most ROIs showed moderate-to-excellent consistency. Power analyses confirmed a sufficient sample size for detecting the reported ICCs. Current modeling suggested that focal-tDCS can induce neurophysiologically relevant electrical field strength in the identified target regions. Hence, we identified accessible, reliable and functionally relevant cortical targets for enhancing novel-word learning. TRR results support the usefulness of the paradigm for future concurrent tDCS–fMRI research. Our study also outlines a general path towards optimization of brain stimulation studies by implementing an empirically informed approach for selecting reliable and relevant target regions and implementation of reliable experimental and imaging paradigms.

## 1 Introduction

A crucial aspect of language acquisition involves learning associations between words and their meanings (Shtyrov, 2012). Because language proficiency is important for social, educational, and vocational success in a globalized world (Greenberg et al., 2001; Kaestle et al., 2001; Young et al., 2002), enhancement of word-learning ability has received substantial attention. In experimental contexts, this has frequently been investigated using associative (Meinzer, 2014; Perceval, 2017; Ramos-Escobar et al., 2021) or contextual (Riemann et al., 2024; Ripollés et al., 2017; Mestres-Missé et al., 2008) novel-word learning paradigms. Those serve as real-world proxies to study (1) the degree and time course of learning under highly controlled conditions, (2) the underlying neural mechanisms, and (3) their potential enhancement by transcranial direct current stimulation (tDCS). The latter is a noninvasive brain stimulation technique with an excellent safety profile (Antal et al., 2026). It allows modulation of cortical excitability and plasticity in targeted brain networks by administering a low-intensity, constant-current stimulus by scalp-attached electrodes (Meinzer et al., 2024).

Some previous studies have indicated that tDCS can enhance novel-word learning in healthy individuals (e.g., Flöel et al., 2008; Meinzer et al., 2014; Perceval et al., 2017, 2020; Riemann et al., 2024). However, outcomes have been variable within and between studies, which has been attributed to individual differences (e.g., baseline learning ability, age, neural network organization) and variations in stimulation protocols (e.g., intensity, focality, timing of tDCS). Moreover, most previous studies on novel-word learning enhancement have stimulated core language regions (e.g., the left temporo-parietal junction, ITPJ, or the left inferior frontal cortex, lIFG), thought to be related to linking novel word-forms and meaning (Rodriguez-Fornells, 2009; Tagarelli et al., 2019). However, neuroimaging research has highlighted that the functional networks involved in novel-word learning can vary depending on task characteristics and the investigated population (e.g., young vs. older adults; Rodriguez-Fornells et al., 2009; Laine & Salmelin, 2009; Shtyrov, 2012). This suggests that stimulation targets need to be optimized for specific experimental paradigms and populations to maximize the effectiveness of tDCS interventions.

Finally, very few brain stimulation studies have formally assessed test-retest reliability (TRR) of experimental paradigms (e.g., Abdelmotaleb et al. 2025), and variability across study participants or repeated sessions may obscure stimulation effects (see Meinzer et al. 2024). This is particularly relevant for studies investigating both behavioral and neural stimulation effects by administering tDCS during functional magnetic resonance imaging (fMRI; Meinzer et al., 2014; Niemann et al., 2024). In this scenario, variability of behavioral responses is compounded by the frequently reported poor reliability of neuroimaging biomarkers (Noble et al., 2019; Elliott et al., 2020). Hence, it is of utmost significance to establish TRR for both behavioral outcomes and task-based markers of brain function, before implementation in tDCS-fMRI studies.

The present study directly addressed these important issues: First, we investigated potential target regions to enhance novel-word learning using an fMRI adaptation of an associative novel word learning paradigm that allows the investigation of the neural mechanisms underlying behavioral performance improvements (Slivinska et al. 2017). This study reported that enhanced proficiency was mainly associated with decreased activity in domain-general and domain-specific regions. We employed a similar paradigm to specifically address the validity of potential target regions for tDCS and to directly investigate brain-behavior correlations (i.e., to determine the functional relevance of potential learning-related changes in brain activity). Second, participants completed two task-related imaging sessions using parallel versions of the experimental paradigm, mimicking crossover designs frequently used to study acute stimulation effects in tDCS-fMRI studies. This aspect aimed to investigate the TRR of behavioral and neuroimaging outcomes, thereby probing its suitability for use in future studies involving tDCS.

## 2 Materials and Methods

### 2.1 Study Overview

The present study was conducted in the context of an ongoing multicenter crossover tDCS-fMRI study, aimed at investigating the behavioral and neural effects of tDCS on learning and memory across functional domains in healthy individuals aged between 18-45 years (Research Unit 5429, https://www.memoslap.de/de/forschung/). The present study reflects the initial phase of the overall project that involved repeated assessment of the different cognitive or motor paradigms used in eight experimental Research Unit (RU) projects (a) to identify potential target regions for active tDCS administration in later project stages and (b) to assess TRR of behavioral and neural outcomes in a repeated measures design. Here, we report results from RU project 3, which investigates potential tDCS effects on language learning ability.

Interested participants were initially screened for eligibility by phone and subsequently completed behavioral and (f)MRI baseline assessments. Afterwards, they participated in four combined tDCS-fMRI sessions. Two sessions involved task-related fMRI and employed an associative picture-pseudoword learning (APPL) task; two additional resting-state fMRI sessions were acquired and will be reported elsewhere. To assess TRR of behavioral and neural outcomes during APPL, we employed placebo (“sham”) tDCS in both sessions. Please note that data acquired in the present study will also serve as a comparator for subsequent active tDCS arms of the RU projects. Hence, we implemented the same experimental setup (e.g., task, electrode configurations) and procedures (e.g., individually optimized electrode positioning, neuronavigated electrode placement; Niemann et al., 2024) that will be used during subsequent study arms involving active tDCS to ensure comparability. The study was pre-registered with the Open Science Framework, and the protocol can be accessed through OSF Registries (https://osf.io/t37u2). **Figure 1A** illustrates the overall study design.

**Figure 1:**
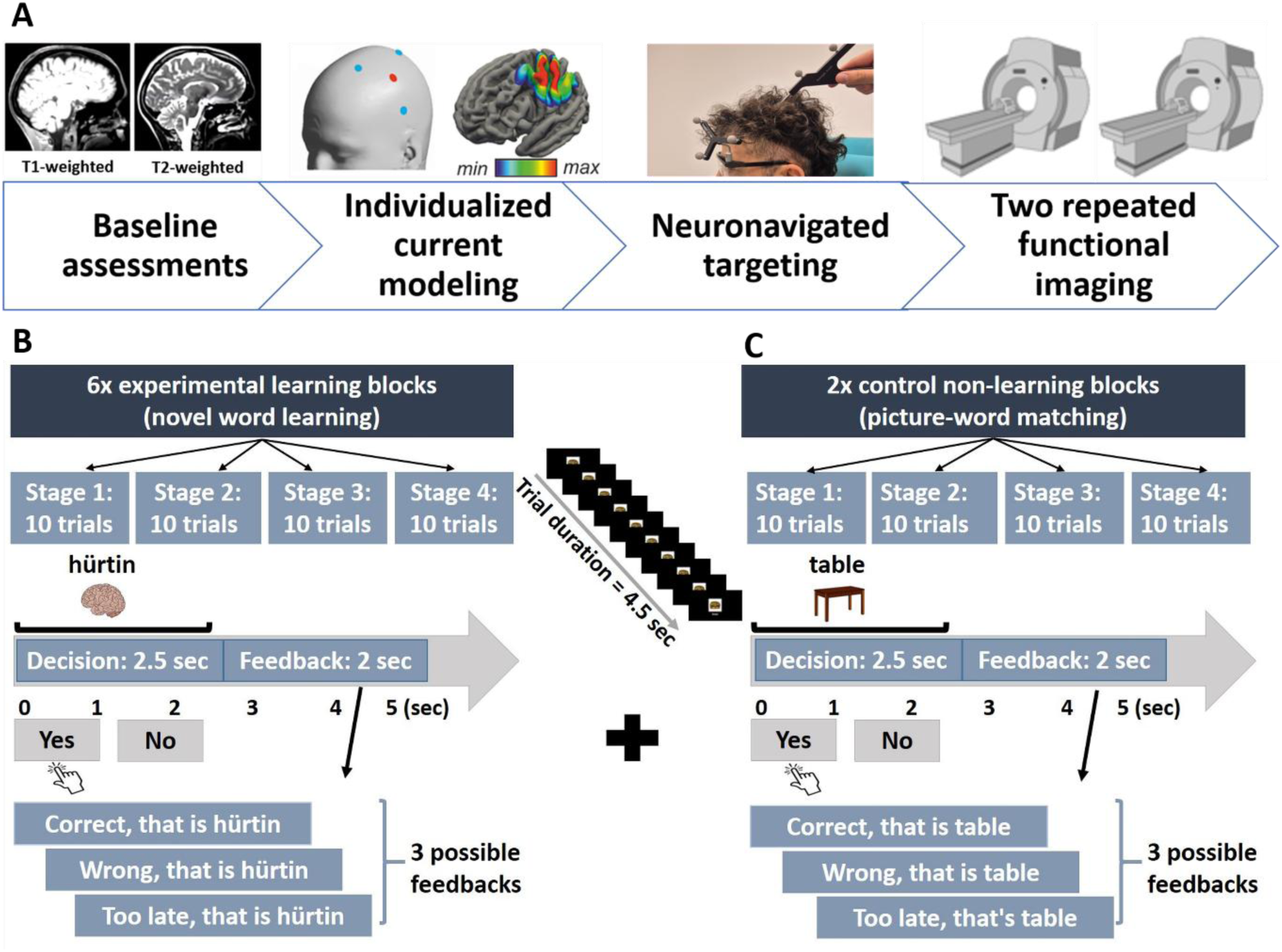
Design and procedure of the intrascanner experimental paradigm; associative picture-pseudoword learning (APPL) task. **Panel A:** Illustrates the overall time flow of the study. Twenty healthy adults completed baseline assessments, including comprehensive neuropsychological testing, a short version of the APPL task, and structural MRI (T1+T2 sequences). Individualized electric-field modeling targeting the left temporoparietal junction (lTPJ) was performed using SimNIBS. This informed neuronavigation-guided focal transcranial direct current stimulation (tDCS) using a 3×1 montage. Each participant completed two task-based fMRI sessions, spaced one week apart. **Panel B and C:** The experiment employed a block design with two identical-structured tasks. Each session comprised six learning blocks involving the APPL task and two blocks of the control task (picture-word matching). Each block comprised four (learning) stages, with 10 trials per stage (5 correct/incorrect pairings). During the APPL task, participants evaluated whether pseudowords were correctly paired with a corresponding object picture and received trial-by-trial feedback. The control task required participants to decide about the correspondence between real-world word-picture pairings. In both tasks, stimuli were presented for 2.5-second (response period), followed by 2-seconds of feedback. Each session consisted of a single fMRI run lasting approximately 29 minutes. To mitigate fatigue and maintain participant engagement, short rest periods were incorporated, including a 4.5 sec. interval between learning stages and an 18 sec. break between blocks. Manual responses (correct/incorrect) were recorded using MRI-compatible response grips. f/MRI = functional magnetic resonance imaging; SimNIBS= Simulation.

### 2.2 Participants

Twenty healthy individuals (10 females and 10 males, mean age = 26; SD = 5.86; range 18-45) participated in this study. All were right handed (Edinburgh Handedness Inventory, Oldfield, 1971) and native German speakers. None reported having any past or present neurological or psychiatric diseases, a history of drug or alcohol abuse, or contraindications for fMRI or tDCS, including use of psychoactive medications, during the initial screening procedures (Antal et al., 2026). The study was approved by the medical ethics committee of the University of Medicine Greifswald (registration number BB015/22). All participants provided written informed consent before study enrollment, and the study was conducted in accordance with the Declaration of Helsinki.

### 2.3 Baseline Assessments

To verify eligibility, all participants completed a comprehensive neuropsychological evaluation to ensure they scored within age-corrected norms and were screened for depression using the German version of the Beck Depression Inventory (BDI-II; Kühner et al., 2007). The following domains were assessed during the neurocognitive assessment: Verbal learning and memory (German Auditory Verbal Learning Test, VLMT, Müller et al., 1997); verbal intelligence and vocabulary knowledge (Multiple-Choice Vocabulary Test, MWT, Lehrl, 2018); executive language functions (Regensburg Verbal Fluency Test, RVFT; Harth & Müller, 2004); visualspatial memory (Rey-Osterrieth Complex **Figure** Test, ROCFT, Zhang et al., 2021); verbal working memory (Digit Span, Woods et al., 2011) and executive functions (Stroop Colour-Word Test, Van der Elst et al., 2006; Trail Making Test A+B, Tombaugh, 2004). All participants scored within age-corrected norms and there was little variability in baseline performance (e.g., mean±SD verbal fluency (RWT) 65.20±.06; vocabulary (MWT) 28.25±2.96; verbal digit span (DS) 16.35±2.43). Nonetheless, we examined whether neuropsychological test performance predicted APPL task accuracy (r=-0.07 to 0.24, none reached significance). Hence, baseline cognitive performance did not robustly predict APPL task performance and was not considered further in behavioral or imaging analyses. Participants also completed a short version of the APPL task. In the present study, this mainly served to familiarize participants with the experimental task before completing it during fMRI. Please note that baseline performance levels on specific tasks have been suggested as a relevant predictor for tDCS response during language and cognitive tasks in health and disease (e.g., Perceval et al., 2020; Riemann et al., 2025; Yucel et al., 2025). Hence, the short version of the APPL task was also conducted to establish a benchmark for classifying age-, sex-, and education-matched participants in the planned active tDCS arms of the RU as high or low performers, to ensure balanced groups.

### 2.4 Task-Related fMRI Paradigm

Each participant completed two task-related fMRI sessions with parallel versions of the APPL paradigm (experimental task), and a control condition (i.e., a lexical decision task, involving object pictures and real words). In each of the two experimental sessions, participants learned 30 unique object picture – pseudoword associations using two parallel sets of stimuli. Each session comprised six learning blocks. Each block comprised four learning stages (10 trials/stage). Learning stages used unique object pictures (N=5) and pseudowords (N=5). Each picture was presented twice per learning stage, once paired with a correct or incorrect pseudoword. Across stages, incorrect pairings were created using different pseudowords. Stimuli were presented using Presentation® software (Version 20.1; Neurobehavioral Systems, Inc., Berkeley, CA, USA) and were projected onto an MRI-compatible screen via a system of mirrors. Behavioral responses were recorded using MRI-compatible response grips (NordicNeuroLab, Norway; www.nordicneurolab.com). FMRI sessions were scheduled at least one week apart. Participants received written and visual instructions detailing the task procedure and practiced use of the response grips prior to each fMRI session. **Figure 1B+C** illustrate the structure of the task-based fMRI sessions and the two tasks.

#### Experimental task

The APPL task was adapted from a previous fMRI study (Sliwinska et al., 2017) and involved learning associations between pictures of real objects and pseudowords across six blocks, each comprising four learning stages. During each block, participants were required to learn associations between objects (pictures of common objects) and legal pseudowords using an explicit instruction- and reinforcement-based approach. The latter was achieved by immediate, trial-specific correct feedback to facilitate learning. The task allows for characterizing overall learning success and the rate of learning over time (performance in learning stages^1-4^ across blocks), while controlling for potentially confounding effects of time on task, novelty of being in the scanner, or fatigue (Sliwinska et al., 2017).

In each learning block, participants were required to identify correct combinations of five object pictures and pseudowords across four learning stages. Pictures were presented twice in each learning stage (10 trials) and were paired with either a correct or incorrect pseudoword. Across learning stages, different pseudowords were used for incorrect combinations, and the same picture was never presented consecutively. Trial duration was 4.5 seconds, and picture-pseudoword pairs were displayed for 2.5 seconds, during which participants were required to indicate the correctness of each stimulus pair by button press using an MRI-compatible response grip with their right hand (index finger: correct; thumb: incorrect). After each trial, written feedback indicated whether the response was correct or incorrect, and the correct picture–pseudoword combination was displayed (2 seconds). If participants did not respond within the 2.5-second time limit, they were notified that they had missed the trial and were still shown the correct association. This ensured that each trial provided accurate feedback for learning. When a picture–pseudoword pair was presented for the first time, participants were instructed to guess and use the feedback provided to facilitate learning. Participants were instructed to respond as quickly and accurately as possible. To ensure a balance between speed and accuracy, participants were informed that only their first response would be recorded.

#### Control task

The control task involved a simple lexical decision task (i.e., matching of object pictures and real words) that mirrored the overall structure of the APPL task (i.e., 40 stimulus pairs per block, 10 per stage). However, to minimize the likelihood of learning or repetition effects, each stage comprised ten unique object pictures (instead of five in the APPL task), paired with five correct or incorrect real words. Across stages, individual pictures were paired once with a correct or incorrect real word.

In **both experimental conditions**, the next trial commenced immediately after the offset of the feedback phase, and the stages were separated by a fixation cross (4.5 seconds). Additionally, a fixation cross was presented for 18 seconds after each task block (i.e., learning or control), and each block began with an information screen (4.5 seconds) that introduced the upcoming experimental condition. The order of the learning and control blocks was randomized but constrained by specific rules (i.e., the first and last blocks were always learning, and control blocks could not be successive).

#### Selection and matching of stimuli

Pictures included 100 color drawings of common objects for the respective parallel versions of the APPL or control tasks (i.e., 2 sessions x 30 or 20 pictures), selected from the Multilingual Picture database (MultiPic, Duñabeitia et al., 2022). This database is the result of an international collaborative effort and features colored images created by a single artist, standardized across multiple languages for name agreement and visual complexity. Pseudowords were created using Wuggy, a pseudoword generator designed explicitly for psycholinguistic research. It supports the creation of pseudowords in multiple languages, including German (https://wuggycode.github.io/wuggy/). Pictures were organized into balanced sets, each paired with a corresponding correct and an incorrect pseudoword. Pairs were randomized and counterbalanced across blocks to avoid systematic biases. For both the APPL and control tasks, parallel sets of images were created and matched based on psycholinguistic and visual parameters, including pseudoword and object picture name length (letters; all pseudowords were bisyllabic), object word frequency (Simple Celex database, http://web.phonetik.uni-frankfurt.de/simplex.html), percentage of modal name usage, and visual object complexity. See **Supplementary Table 1** for details and statistical comparison of the respective stimulus sets.

### 2.5 Focal Sham tDCS

Sham tDCS was administered with an MRI-compatible multichannel direct current stimulator and equipment (DC-STIMULATOR MC MR, NeuroConn GmbH, Germany) and a 3×1 focal setup involving a central anode and three equidistantly arranged cathodes (electrode diameter: 2 cm; anode-cathode center distance: 4 cm). The arrangement of the three cathodes was standardized using a 3D-printed thermoplastic spacer (Niemann et al., 2024), ensuring a consistent anode-cathode center-to-center distance and equal spacing between the three cathodes. Additional procedures were in place to ensure comparability of procedures with subsequently planned active tDCS arm: i.e., individualized current modeling (using structural imaging data acquired during the baseline scan) was performed for each participant to optimize theoretical current flow to the envisaged target region, and electrode placement on the scalp was guided by neuronavigation (Niemann et al., 2024). In the present study, which did not involve active stimulation, the center anode was placed over the lTPJ. This decision was based on previous behavioral studies suggesting beneficial effects of lTPJ tDCS on novel word-learning (Flöel et al., 2008; Meinzer et al., 2014; Perceval et al., 2017; Filippova et al., 2023; Kurmakaeva et al., 2023). The sham protocol comprised a 10-second ramp-up to 2 mA at the start of the stimulation, which remained constant for 20 seconds before ramping down over 10-seconds. The procedure mimics the initial physical sensation elicited by prolonged active tDCS to ensure participant blinding, without inducing neurophysiological effects. Sham tDCS was completed immediately prior to initiating the respective functional imaging sessions.

### 2.6 FMRI Data Acquisition

MRI data were acquired using a 3.0 Tesla Siemens MAGNETOM Vida scanner (Siemens Healthineers, Germany) at University Medicine Greifswald, equipped with a 64-channel head-neck coil. Functional MRI data were collected using a multiband echo-planar imaging (EPI) sequence developed by the Center for Magnetic Resonance Research at the University of Minnesota (https://www.cmrr.umn.edu/multiband/). This sequence incorporated simultaneous multislice (SMS) acceleration and was optimized for blood oxygenation level-dependent (BOLD) contrast to achieve high temporal resolution while minimizing slice repetition time and maximizing whole-brain coverage (Setsompop et al., 2012; Xu et al., 2013). Task-related fMRI scans were acquired with the following parameters: 110 × 110 matrix, spatial resolution of 2 × 2 mm, slice thickness of 2 mm (no interslice gap), repetition time (TR) = 1000 ms, echo time (TE) = 30 ms, flip angle = 60°, field of view (FOV) = 220 mm, and a multiband acceleration factor of 6. Each session lasted approximately 29 minutes, during which 1760 volumes were collected with phase encoding in the anterior-to-posterior (AP) direction. To minimize image distortions and signal drop-out, the protocol included an echo spacing of 620 microseconds and parallel imaging using GeneRalized Autocalibrating Partially Parallel Acquisitions (Grappa) with an acceleration factor of 2. Slice timing was interleaved to reduce motion-related artifacts, and field maps were acquired for subsequent geometric distortion correction. We also acquired high-resolution T1-weighted images (0.9 × 0.9 × 0.9 mm³, TR = 2700 ms, TE = 3.7 ms, inversion time = 1090 ms, flip angle = 9°).

### 2.7 Statistical Analysis

#### 2.7.1 Behavioral Data Analysis

Performance in the APPL task was assessed using both response accuracy and reaction time (RT) for correct responses. Generalized linear mixed-effects models (GLMMs) were used to examine the effects of task condition (learning vs. control), stage (1–4), and their interaction. Each stage (1–4) within a block used different sets of stimuli, ensuring no repetition across stages. All models included task session (1 -2), task version (set1 – set2), and block (learning: N=6; control: N=2) as fixed effects. To account for inter-individual variability, subject-specific random intercepts were included. The random-effects structure was determined by balancing model complexity and estimation stability. More complex structures (e.g., random slopes) were not included due to convergence issues and risk of overparameterization given the sample size and design. All analyses were conducted using the glmer function from the lme4 and lmerTest packages in R (version 4.4.1).

For accuracy, a binomial GLMM with a logit link was used to model correct vs. incorrect responses. For RT, a GLMM with an inverse Gaussian distribution and log links was fitted to account for the skew in RT data (Lo & Sally Andrews, 2015). RT analyses were based on correct trials only. Post hoc comparisons of estimated marginal means (EMMs) were conducted using the *emmeans* package to visualize interactions between task condition and stage. EMMs were computed for each combination of task condition and stage to visualize how accuracy and RT varied between experimental conditions across stages.

### 2.8 fMRI Data Analysis

#### 2.8.1 Preprocessing

MATLAB (R2021b) and statistical parametric mapping software (SPM12, v7771, http://fil.ion.ucl.ac.uk/spm/) were used to analyze fMRI data. Skull stripping, bias correction, and normalization to the Montreal Neurological Institute 152 template were applied to structural T1 images while maintaining the original voxel size. Slice timing correction (TR = 1s, 72 slices), realignment, and unwarping were applied during preprocessing. T1 and functional images were coregistered, and deformation fields obtained from T1 segmentation were used to normalize functional images to MNI space. Finally, to enhance the signal-to-noise ratio and satisfy GLM criteria, spatial smoothing with a 6 mm FWHM Gaussian kernel was applied (Kuhnke, P. GitHub: https://github.com/PhilKuhnke, 25.09.2025).

We modeled participant-specific functional data using a first-level GLM containing regressors for learning and control conditions across all four stages. Each event was defined as a 2.5s fixed-duration boxcar, reflecting the actual trial length, and convolved with the canonical hemodynamic response function (HRF). All event timings were synchronized to the functional run onset via behavioral log files.

Data from the two task-fMRI sessions were concatenated into a single design matrix. To account for head motion artifacts, nuisance regressors were included: six motion parameters from image realignment and framewise displacement. The following contrasts of interest were generated for each study participant: (1) The comparison of APPL *>* control tasks and the inverse contrast was used to directly compare overall activity patterns elicited by the two experimental conditions. (2) We also generated task-specific contrasts (i.e., APPL or control > implicit baseline). The latter were used in subsequent a priori region-of-interest (ROI, see below for details) analyses to explore the time course of activity in both tasks across the four stages and to calculate correlations with behavioral performance.

Subsequently, the following group-level analyses were performed:

1. We initially conducted a second-level whole-brain analysis to examine overall task-related activation differences between the APPL paradigm and the control condition. A voxel-wise statistical threshold of p < 0.001 (uncorrected) was applied for initial cluster formation, with a family-wise error (FWE) corrected cluster level threshold of p < 0.05 (cluster extent k=192). Anatomical labeling of significant clusters used the Harvard-Oxford cortical and subcortical structural atlases (https://fsl.fmrib.ox.ac.uk/fsl/fslwiki/). Activation maps were overlaid on an MNI152 template and visualized using MRIcroGL (http://www.mricro.com).
2. To investigate potential (a) **changes in activity across learning stages** and (b) the relationship **between behavioral performance and brain activity changes during the APPL task**, we conducted two a priori ROI-based analyses. In line with current guidelines for unbiased ROI specification (Poldrack, 2007; Tong et al., 2016), ROIs were selected based on results from a recent neuroanatomical meta-analysis of lexical learning (Tagarelli et al., 2019). This study identified a consistent network of regions involved in lexical learning, including the lIFG (peak MNI coordinates x/y/z left pars opercularis -46/6/28; pars triangularis -46/28/20), lTPJ (-60,/-43/22), left anterior insula (lAnIns: -32/24/2), left superior parietal lobule (lSPL: -32/-60/54), left supplementary motor area (lSMA: -4/6/60), left fusiform gyrus (lFuG: -40/-54/-18) and right anterior insula (rAnIns: 2/22/0). These regions are broadly implicated in phonological processing, semantic integration, and procedural learning mechanisms (Rodríguez-Fornells et al., 2009), supporting their relevance for encoding and retrieval processes involved in novel word learning. For the ROI analyses, spherical ROIs with a radius of 5 mm were created, centered on the above regions. Mean beta values (parameter estimates) were extracted for each subject, stage, task, and ROI using the MarsBaR toolbox (http://marsbar.sourceforge.net/; Brett et al., 2002). Only voxels exceeding a percentile-based activation threshold, corresponding to the top 25% voxels in each ROI, were used (Tong et al., 2016). This allowed the investigation of:

a. Changes of activity across stages for the APPL and control tasks.
b. Associations between changes in brain activity during the APPL task across learning stages and behavioral outcomes (accuracy, response latency). This analysis involved a repeated-measures correlation analysis using the *rmcorr* package in R (https://cran.r-project.org/web/packages/rmcorr/; Bakdash & Marusich, 2017). The approach was chosen because it appropriately accounts for dependency within-subject repeated measures, allowing for robust estimation of consistent individual-level, brain–behavior associations across multiple time points. This method estimates the common within-subject association between neural activity and behavioral performance across repeated observations, while accounting for the non-independence of measurements within individuals. Specifically, rmcorr controls for between-subject differences in baseline levels by fitting parallel regression lines for each participant, thereby isolating the shared within-subject relationship.

### 2.9. Test-retest-reliability analyses

To evaluate the TRR of both behavioral performance and fMRI activation across the two sessions that used parallel stimulus sets, we performed an ICC analysis (Caceres et al., 2009). The ICC is a statistical measure that quantifies the degree of consistency or agreement across repeated measurements, making it suitable for assessing TRR. It accounts for both within- and between-subject variability, thereby providing a comprehensive estimate of measurement stability over time (Weir, 2005; Koo & Li, 2016). ICC values were interpreted according to established benchmarks (Koo & Li, 2016): < 0.5 indicated poor reliability, 0.5–0.75 moderate reliability, 0.75–0.9 good reliability, and values > 0.9 excellent reliability.

**Behavioral TRR** considered performance accuracy and RT across learning stages for each subject in both task sessions (sessions 1+2). ICC estimates and their 95% confidence intervals were computed using the *irr* package in R, applying a two-way mixed-effects model with absolute agreement for single measurements (ICC [3,1]), following McGraw and Wong (1996). This model accounts for both systematic and random sources of measurement variability, enabling assessment of absolute agreement between sessions at the individual level.

To investigate **fMRI reliability**, we employed the Python-based PyReliMRI tool (Demidenko et al., 2024) to compute ICCs based on activity (beta values) obtained in the two fMRI sessions using data from the contrast learning > baseline. The implicit baseline was chosen to capture overall task-related activation associated with novel-word learning (Note: ICCs based on the complex contrast learning vs. control task would have been influenced by variability in either task. Hence, we decided to report ICCs only for the task of interest).

We describe four analyses: (1) A whole brain analysis, aimed at providing an illustrative overview of the voxel-wise ICC distributing in our data. (2) A ROI analysis providing a formal evaluation and description of voxel-wise ICCs within the task-active regions. Two additional ROI analyses tested (3) ICCs in the above-described a priori ROIs and (4) ROIs centered around peak ICC coordinates obtained from the voxel-wise ICC map within clusters of task-related activation. These ROIs represent voxels exhibiting the highest local reliability within the task-relevant network. Additional information is provided in the **Supplementary Methods**.

We also performed a *formal power analysis* to quantify the adequacy of our sample size for detecting the observed ICC effects in a priori ROIs and ROIs centered around peak ICC values [i.e., analyses (3)+(4) described above]. Power was estimated using the non-central F approximation for the ICC F-test under a two-way mixed model with k=2 sessions and α=0.05 (Donner & Eliasziw, 1987; Walter, Eliasziw & Donner, 1998).

### 2.10. Post-hoc Current Modeling

To address targeting of brain regions identified by fMRI with focal tDCS, we conducted post-hoc individualized current modeling based on structural imaging of our sample (Niemann et al. 2024; details of the methods are provided in the Supplementary Methods). In brief, this analysis aimed to illustrate current flow induced by focal 3×1 tDCS montages targeting three selected functionally relevant brain regions for novel word learning identified in the fMRI analyses (i.e., left IFG, TPJ, SMA). Outcomes are provided at the whole brain level, to illustrate the extent and distribution of the simulated current flow relative to the respective target regions in our sample. To directly quantify current intensity in each region, we also extracted field strength from ROIs centered on the respective target coordinates.

## 3. Results

### 3.1. Behavioral Data Analysis

Overall, we confirmed effective acquisition of the novel vocabulary, as illustrated by a gradual increase in accuracy and a decrease in RT for correct responses across the four learning stages. Participants in the learning condition reached asymptotic performance levels at the third stage **(see Fig. 2).** As expected, accuracy levels were close to ceiling for the control task. **Figure 2** illustrates the behavioral results. In the following, only the most relevant effects are described; please see **Supplementary Table 2** for the full model.

**Figure 2:**
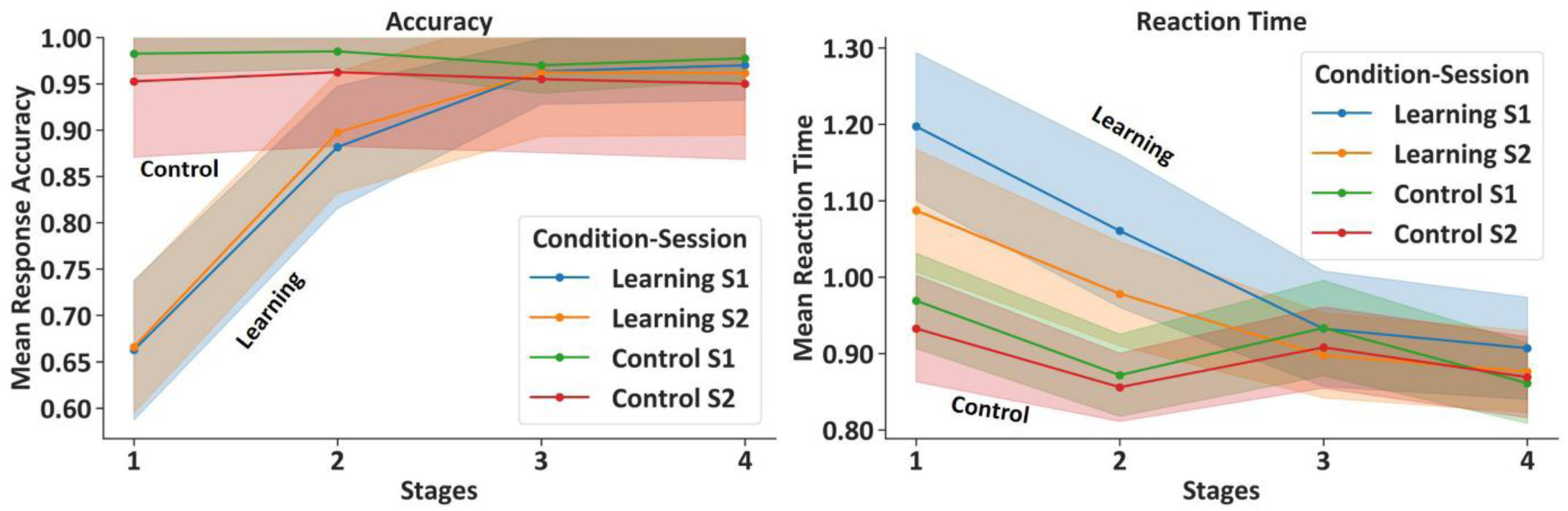
Performance accuracy and reaction time (RT) for associative picture-pseudoword learning (APPL) *and control tasks*. **Left panel:** Mean response accuracy and **right panel:** mean RT (seconds) across four learning stages (1–4) for the *APPL task* and the *control task.* Results are shown separately for the *first fMRI session* and the *second fMRI session*, with data averaged across 20 participants. Shaded areas represent standard deviation. S= task session; fMRI= functional magnetic resonance imaging.

#### 3.1.1. Response Accuracy

The GLMM analysis revealed a significant main effect of task (APPL vs. control; estimate = -4.05, SE = 0.26, z = -15.44, p <0.001, 95% CI = [-4.55, -3.55]) and a significant interaction between task and stage (estimate = 1.21, SE = 0.10, z = 12.40, p <0.001, 95% CI = [1.01, 1.41]). Post-hoc pairwise comparisons with estimated marginal means (EMMs) demonstrated better performance for the control compared to the APPL task in the first two stages (stage 1: estimate = 1.32, SE = 0.12, z = 10.77, p < 0.001; stage 2: estimate = 0.61, SE = 0.10, z = 6.21, p < 0.001), while performance levels at stage three were comparable (estimate = -0.09, SE = 0.11, z = -0.82, p = 0.41). At stage four, accuracy was higher for the APPL task (estimate = -0.76, SE = 0.15, z = -5.07, p < 0.001). Furthermore, task version had a small but significant effect on accuracy (estimate = 0.21, SE = 0.06, z = 3.29, p =0.001), with higher overall accuracy for task version 2. While significant, the task version effect is numerically small (mean/SD Set1: 0.886±0.317, Set 2: 0.903±0.296) and likely reflects minor differences between the two stimulus sets. There was no significant effect of session (p =0.486).

#### 3.1.2. Reaction Time (RT)

Overall, RT were significantly slower in the APPL task compared to the control condition (estimate = 0.220, SE = 0.010, t = 22.09, p <0.001, 95% CI [0.200, 0.240]). Additionally, there was a significant interaction between task and stage (estimate = -0.057, SE = 0.004, t = -16.15, p <0.001, 95% CI [-0.065, -0.050]), reflecting greater RT decreases across stages during the APPL task. Post-hoc tests showed that the most significant difference between the tasks was found at stage 1 (estimate = 0.156, SE = 0.007, z = 21.46, p < 0.0001). Participants were significantly faster in the second experimental session (estimate = -0.038, SE = 0.004, t = -10.70, *p* < 0.001, 95% CI [-0.045, -0.031]), but task version (version 2 vs. 1) did not significantly affect RT (estimate = -0.001, SE = 0.004, t = -0.28, *p* = .784, 95% CI [-0.008, 0.006]).

### 3.2. Functional Imaging Results

#### 3.2.1. Whole-Brain fMRI Analysis

Clusters of significant activity elicited by the respective tasks (learning, control) are illustrated in **Figure 3A+B.** Across blocks, the most pronounced activity during the learning task was found in bilateral occipito-temporal, lateral and medial frontal and temporo-parietal regions, a pattern consistent with the task requirements. At the same statistical threshold, activity elicited by the control task was most pronounced in occipito-temporal regions.

**Figure 3:**
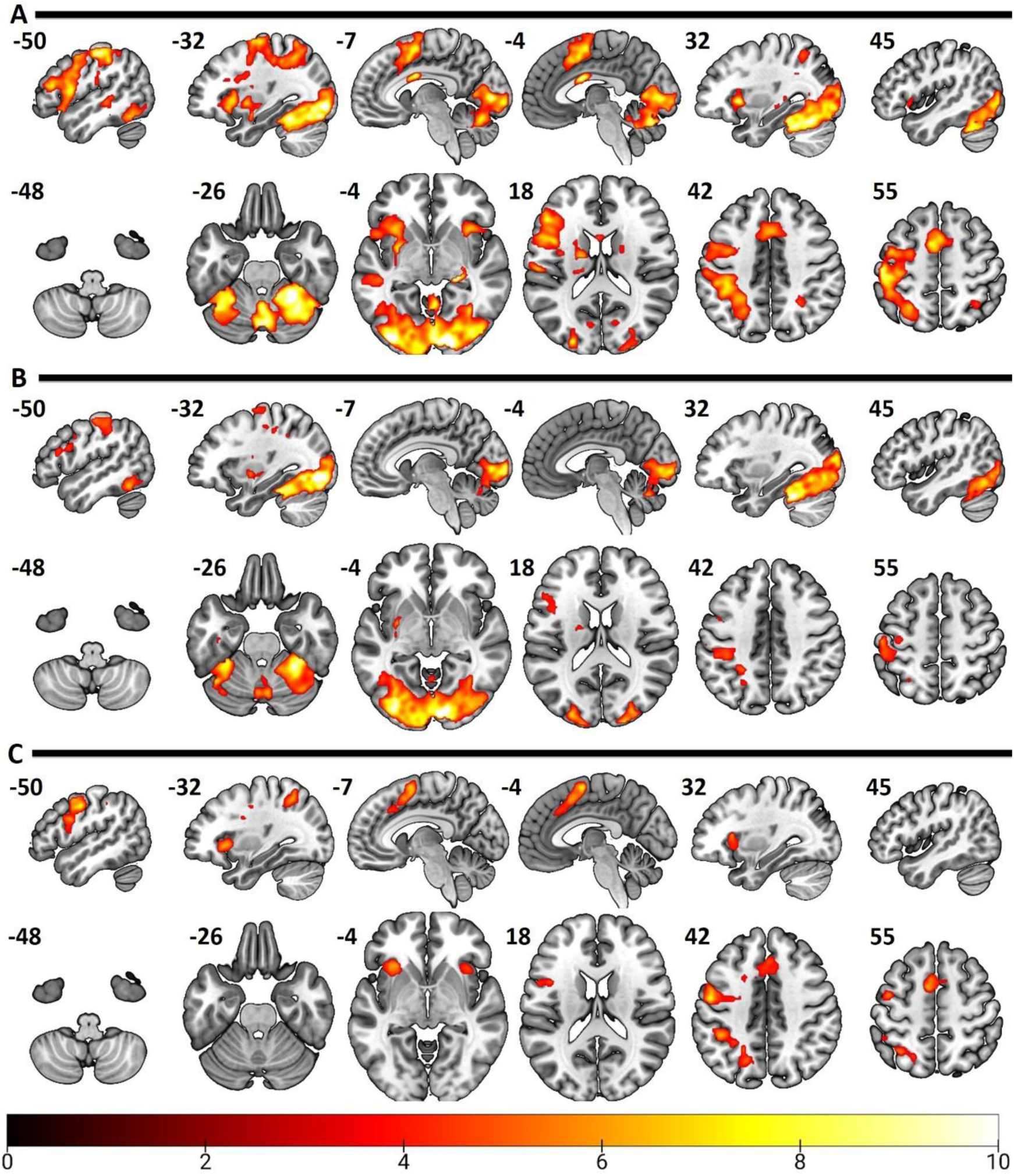
Activity patterns associated with (A) APPL task > implicit baseline, (B) control > implicit baseline, (C) APPL > control. Brain activation patterns are shown for sagittal and axial planes. Statistical parametric maps are overlaid on a standard MNI brain template. Only significant clusters derived from the group-level analyses across both task sessions for all 20 participants are shown. A cluster-level family-wise error (FWE) correction of p< 0.05 was applied, with a voxel-level significance threshold set to *p* < 0.001. The color bar indicates values of the T-statistic. APPL= associative picture-pseudoword learning; MNI**=** Montreal Neurological Institute.

The direct comparison of the two tasks revealed significantly greater activity during the APPL task in 10 clusters **(Figure 3C)**. The strongest effects were found bilaterally in the insula, pars opercularis and pars triangularis of lIFG, lSMA, and bilateral premotor regions. No significant differences were found for the inverse contrast (control > APPL task). See **Supplementary Table 3** for details.

#### 3.2.2. Changes in Task-Related Activity Across Stages

Analysis of brain activity across learning stages revealed marked differences between the APPL (learning) and control tasks **(see Figure 4)**. APPL-related activity (relative to the implicit baseline) decreased in all a priori ROIs across stages, including the IIFG, IAnIns, rAnIns, lSMA, and ISPL. This decreasing pattern suggests that these regions are particularly engaged during the early phases of learning, likely reflecting processes related to effortful encoding. In contrast, activation in the control task in these regions was less pronounced and remained relatively stable across stages. Activity during the APPL task also decreased in the lTPJ and lFuG, though the differences between task conditions were less pronounced.

**Figure 4:**
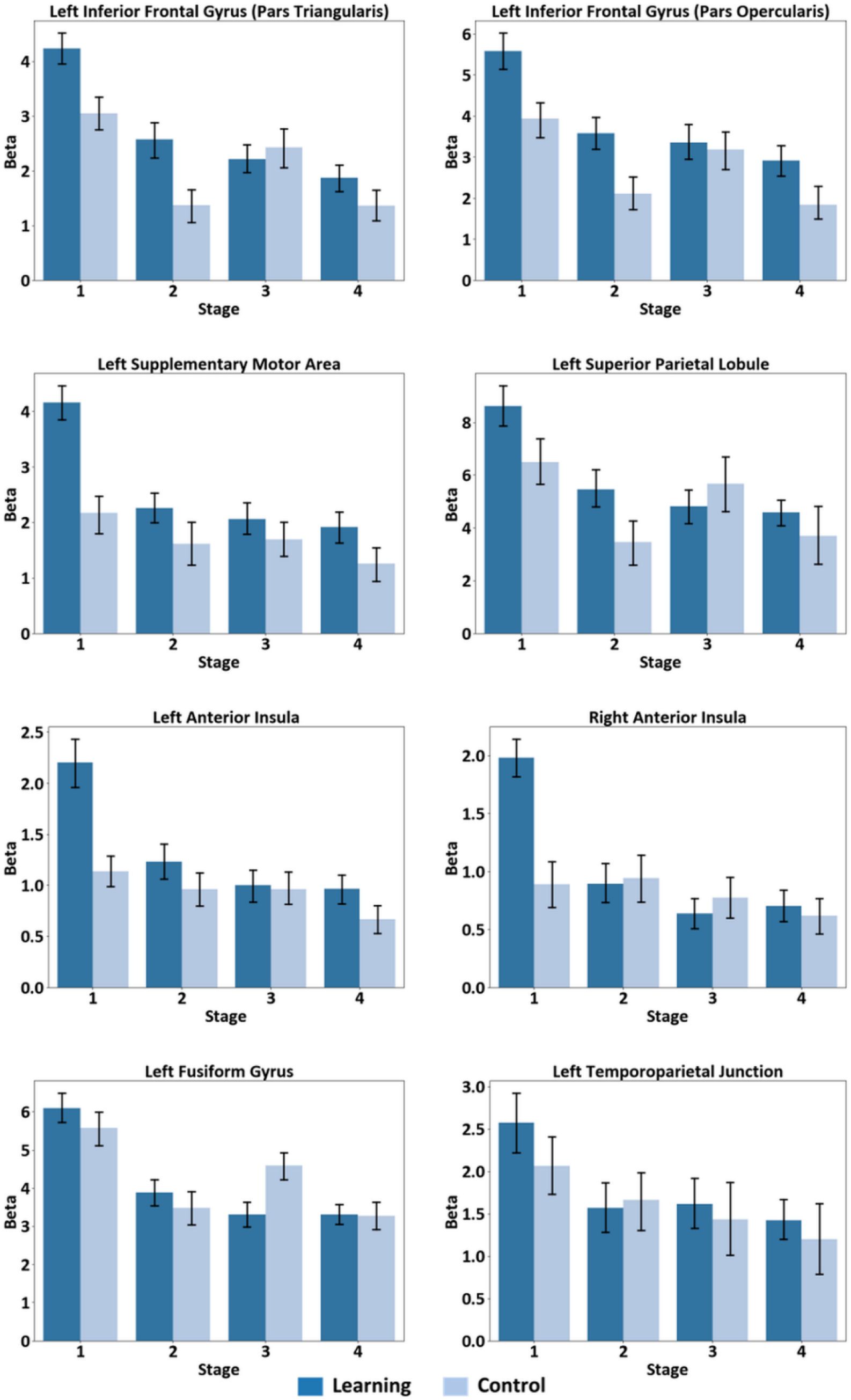
*Task-related brain activity changes across the four stages in* a priori *regions-of-interests (ROIs).* Bar plots show mean beta values for the learning (APPL, dark blue) and control (light blue) tasks across four stages (1–4), pooled across the two imaging sessions. Within each region, only the top 25% most active voxels were used to compute mean beta values. Error bars represent the standard error of the mean. **APPL: associative picture-pseudoword learning, lIFGpt: Left Inferior Frontal Gyrus (Pars Triangularis), lIFGpo: Left Inferior Frontal Gyrus (Pars Opercularis), lSMA: Left Supplementary Motor Area, lSPL: Left Superior Parietal Lobule, lAnIns: Left Anterior Insula, rAnIns: Right Anterior Insula, lFuG: Left Fusiform Gyrus, lTPJ: Left Temporoparietal Junction.**

### **3.3.** Brain-Behavior Relationships

#### 3.3.1. Learning (APPL task)

The a priori ROI analysis revealed consistent associations between task-related neural activity and learning-related behavioral improvement. Specifically, reduced task-related activity in all a priori ROIs was significantly associated with (a) increased accuracy and (b) reduced RT across learning stages (see **Figure 5** and **Supplementary Figure 1)**. Specifically, faster RT were associated with decreased activation in all ROIs, with correlation coefficients ranging from r = 0.45 to 0.71. Correlations between increased accuracy and reduced task-related activity ranged from r = -0.51 to -0.77. The strongest associations for RT were observed for the left fusiform gyrus (r = 0.71) and left IFG (r = 0.69), while accuracy showed the strongest correlation with the left fusiform gyrus (r = -0.77).

**Figure 5:**
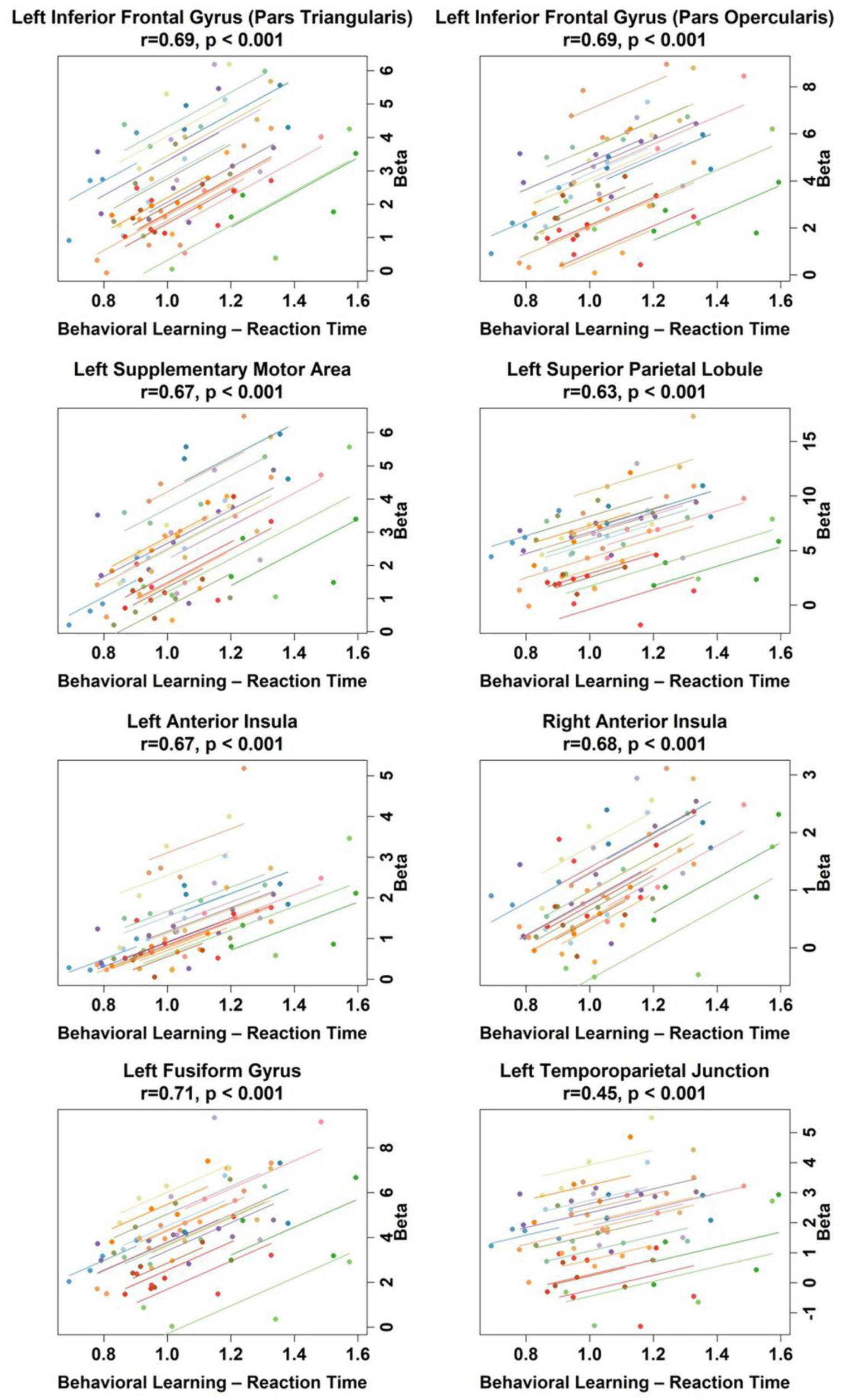
Repeated-measures correlations between behavioral learning performance (reaction time, RT) and brain activity (beta values) across learning stages in a priori regions-of-interest (ROIs). Points represent participants and stages (20 participants, 4 stages), pooled across the two imaging sessions. Lines represent individual participant-specific trends across the four stages (n = 20), illustrating within-subject trajectories. Shorter RT (faster response latency) was associated with lower beta values. Correlation coefficients (r) and associated p-values are reported for each ROI. Note: The y-axis is plotted on the right to emphasize the decrease in RT across the four stages. **lIFGpt: Left Inferior Frontal Gyrus (Pars Triangularis), lIFGpo: Left Inferior Frontal Gyrus (Pars Opercularis), lSMA: Left Supplementary Motor Area, lSPL: Left Superior Parietal Lobule, lAnIns: Left Anterior Insula, rAnIns: Right Anterior Insula, lFuG: Left Fusiform Gyrus, lTPJ: Left Temporoparietal Junction.**

#### 3.3.2. Control Task

Decreased RT in the control task was also associated with reduced activity in six of the eight a priori ROIs, including the two subportions of the IIFG and IAnIns, lFuG, ISPL, and lSMA. No correlations between RT and activity changes were observed in the IAnIns and rAnIns **(Supplementary Figure 2)**. Overall, brain-behavior correlations were weaker compared to those observed for the learning task and ranged from r=0.5 to 0.51. No significant correlations were observed for response accuracy (r=-0.07 to -0.29; **Supplementary Figure 3)**.

### 3.4. Test-Retest Reliability (TRR)

#### 3.4.1. Behavioral Outcomes

Excellent reliability was found for response accuracy (ICC = 0.909 (95% CI [0.861, 0.94]), suggesting comparable performance across repeated sessions. The F-test associated with the ICC was statistically significant (F (79, 78.7) = 21.2, p < 0.001), indicating that the observed level of agreement is highly unlikely to have occurred by chance. Overall, RT also showed good consistency between sessions (ICC = 0.811 (95% CI [0.429, 0.917]), although TRR was lower compared to response accuracy. The ICC F-test for RT was also statistically significant (F (79, 6.75) = 15.3, p = 0.0006). These results highlight the reliability of the APPL task used in this study and suggest stable performance over time.

#### 3.4.2. Imaging Outcomes

ICCs illustrating the degree of consistency of brain activity in the two APPL sessions are illustrated for the entire brain (Figure 6A) and also for regions with significant activity during the APPL task **(Figure 6B)**. The former analysis showed highly variable ICCs across the entire brain, with higher ICCs mainly in task-relevant brain regions, including lateral and medial frontal, temporo-parietal, and occipital regions in the left hemisphere. The highest ICCs were found in the lIFG (MNI coordinates x/y/z pars triangularis: (-45/41/6; ICC = 0.892, 95% CI [0.749, 0.956]; pars opercularis -46/6/24; ICC = 0.908, 95% CI [0.784, 0.962]), lSPL (MNI: -36/-52/50; ICC = 0.905, 95% CI [0.653, 0.936]), lSMA (MNI: - 55/-36/33; ICC = 0.848, 95% CI [0.657, 0.937]), and lTPJ (MNI: -59/-38/24; ICC = 0.854, 95% CI [0.651, 0.935]). Approximately 81% of task-related activity showed moderate-to-excellent TRR (median ICC= 0.696; for details see **Supplementary Figure 4)**, highlighting consistent neural signatures within core regions of the functional network associated with novel-word learning and potential targets for noninvasive brain stimulation.

**Figure 6:**
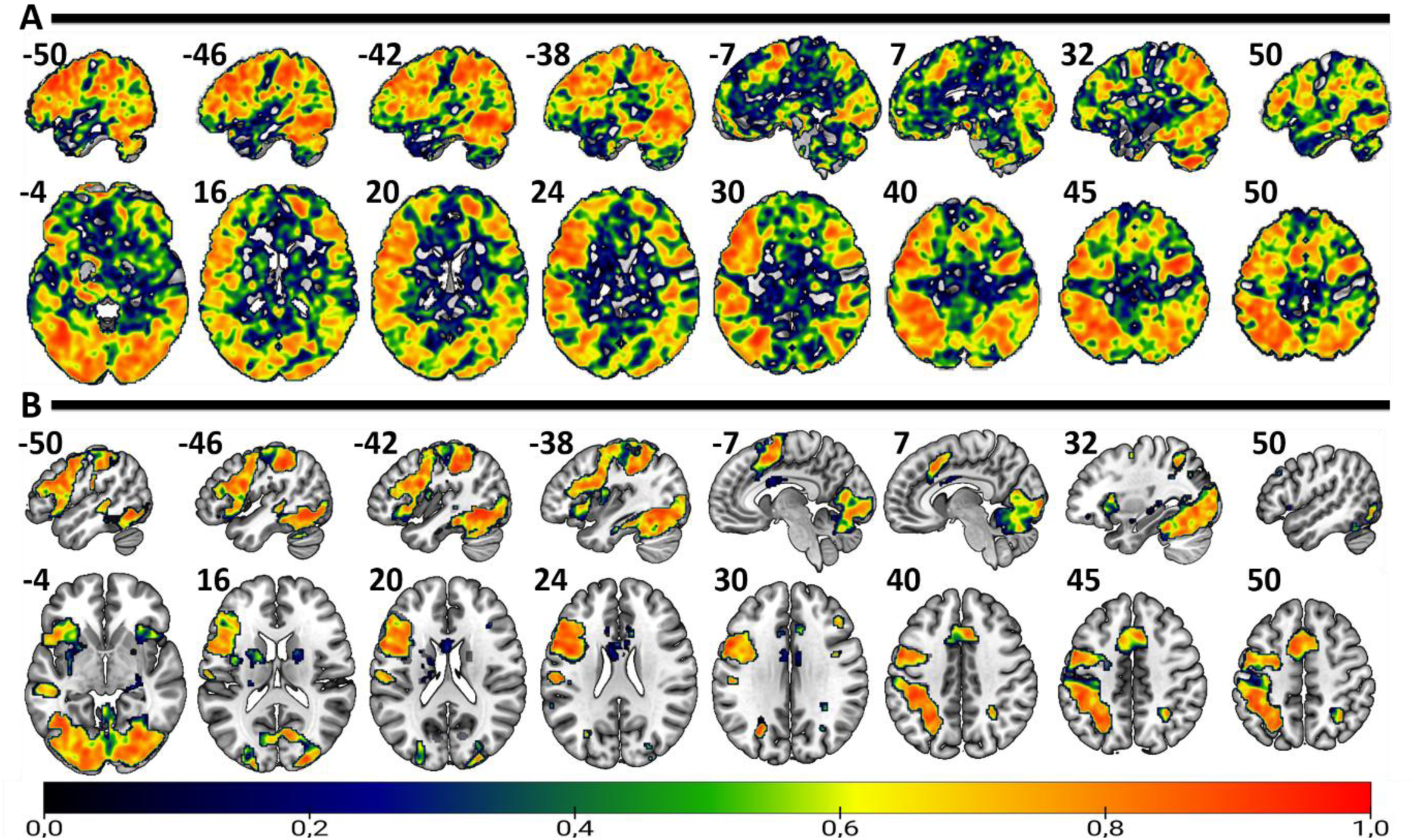
Intraclass correlation coefficient (ICC) maps illustrating the reliability of brain activations during the APPL task across two sessions. The figure displays ICCs for the whole brain (A) and for regions active during the APPL task (B), overlaid on an MNI template, showing the distribution of ICC values. The color bar ranges from 0 to 1, with red representing high reliability and blue indicating low reliability. **APPL=associative picture-pseudoword learning; MNI=**Montreal Neurological Institute.

Six of the eight **a priori ROIs** showed good-to-excellent reliability (ICC ≥ 0.60), with the left SMA (ICC = 0.817), left IFG pars opercularis (ICC = 0.785), and left SPL (ICC = 0.763) demonstrating the highest consistency across sessions. The left IFG pars triangularis showed moderate reliability (ICC = 0.543), while the right anterior insula showed poor reliability (ICC = 0.390). The mean ICC across all a priori ROIs was 0.667 (SD = 0.144). All five **ICC-peak ROIs** demonstrated good-to-excellent reliability (ICC range: 0.796–0.879). The mean ICC across ICC-peak ROIs was 0.840 (SD = 0.039). Notably, the left IFG (both pars opercularis and pars triangularis at the ICC-peak coordinates) showed the highest reliability (ICC = 0.879), consistent with this region’s role as a core processing node for novel-word learning. Please see **Supplementary Figures 5+6** for additional information.

#### 3.4.3. ICC power analyses

Seven of eight **a priori ROIs** achieved power ≥0.80 with the current sample size of N=20, indicating that the study was adequately powered to detect the observed ICC effects. The only exception was the right anterior insula (rAnIns; power = 0.586), which would require N ≈ 35 participants to achieve 80% power. All five **ICC-peak ROIs** were highly powered (achieved power = 1.000), requiring only N = 5 participants for 80% power. For details, see **Supplementary Tables 4+5** and **Supplementary Figures 5+6**.

### 3.5. Post-hoc current modeling

Current modeling showed excellent correspondence of peak current flow relative to the target regions (illustrated in **Figure 7**). This was confirmed by the ROI analyses demonstrating that neurophysiologically relevant field strengths (>0.2 V/m; Thielscher et al. 2026) are achieved in all target regions (mean |E|±SD (V/m) lIFG: 0.21±0.05; lTPJ: 0.28±0.08; lSMA: 0.33±0.68; for details see Supplementary Methods). For the lSMA, relatively high electric field intensities also extend beyond the target region and this region showed the highest inter-individual variability, likely due to current shunting via cerebrospinal fluid in the interhemispheric fissure.

**Figure 7:**
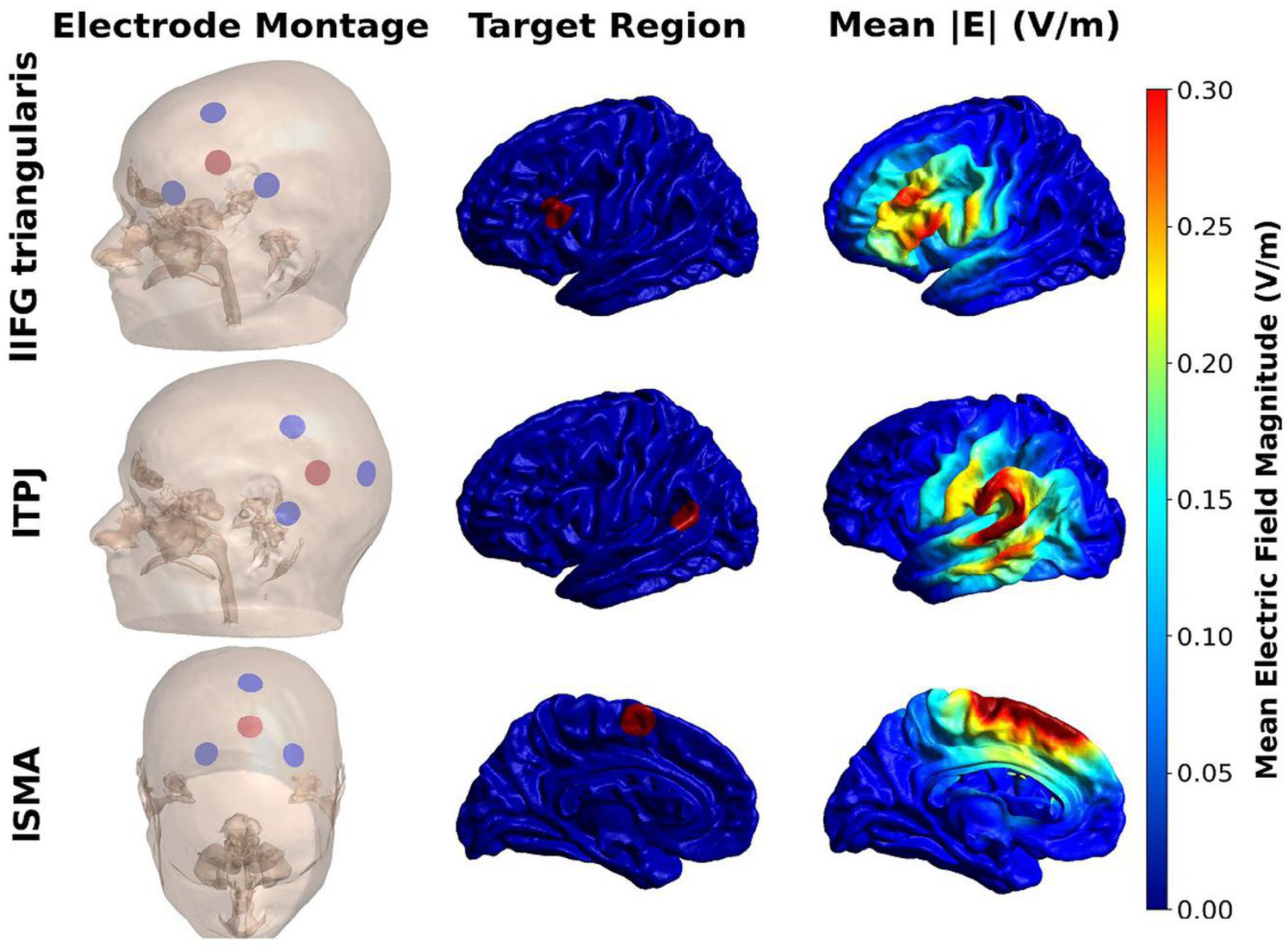
Group-averaged electric field distributions induced by focal tDCS targeting three of the identified functionally relevant cortical regions for the APPL task. Individual electric field simulations (n=20) were performed for three 3×1 montage targeting the left inferior frontal gyrus (lIFG), left temporo-parietal junction (lTPJ) and left supplementary motor cortex (lSMA). **Left column:** Electrode configurations showing the location of the anode (red) and the three surrounding cathodes (blue) on a representative subject’s head model in native space. **Middle column:** 10 mm spherical regions of interest (ROIs) centered on each target coordinate, projected onto the FsAverage cortical surface to illustrate anatomical location of the intended target regions. **Right column:** Group-averaged electric field magnitude (|E|, V/m) on the FsAverage pial surface (common color scale: 0–0.3 V/m). **Rows** represent three montages: lIFG pars triangularis (top), lTPJ (middle), and lSMA (bottom).

## 4. Discussion

This study aimed to (I) identify potential task-specific cortical target regions for enhancing novel word learning with tDCS, and (ii) to evaluate the TRR of both behavioral and neural outcomes across repeated sessions using a novel fMRI adaptation of the APPL task. This yielded the following main outcomes: (1) Participants successfully acquired novel word forms, indexed by a gradual increase in accuracy and reduced RT across the learning stages. Behavioral outcomes also showed good-to-excellent consistency across sessions. Notably, accuracy showed ceiling effects in the later learning stages, indicating limited sensitivity to detect further improvements. While this confirms successful acquisition of the novel vocabulary, accuracy measures may be less suitable for capturing the effects of NIBS (e.g., tDCS). RT provides a more sensitive index of learning dynamics across stages and is suggested as primary behavioral outcome for future implementations of the current APPL paradigm, particularly in the context of tDCS–fMRI studies. (2) Robust task-related activity was found in regions implicated in language processing and learning, as well as domain-general regions. The identified accessible and functionally relevant candidate target regions for tDCS included the lIFG, ISMA, and ISPL. (3) APPL-related activity decreased across learning stages, which was correlated with learning success. This overall pattern suggests improved efficiency in retrieving novel word-forms and consolidation of the acquired object-picture and non-word associations. The functional relevance and specificity of this pattern to novel-word learning is emphasized by the analysis of the control task data, which showed no consistent pattern of activity changes across stages. (4) Overall, the identified functionally relevant regions showed moderate-to-excellent TRR in the whole-brain and ROI analyses, thereby highlighting their roles as core nodes for acquiring novel word meanings. This was further emphasized by a formal power analysis, which suggested that our sample size was sufficient for detecting the reported ICCs. (5) Finally, post-hoc current modeling for selected target regions confirmed that focal tDCS can induce neurophysiologically relevant electrical field strengths in the identified target regions. In sum, we identified several accessible, functionally relevant, and reliable cortical target regions for enhancing novel word learning, and the overall results of the TRR analysis support the utility of the new APPL task for future use in concurrent tDCS–fMRI studies.

The experimental task activated a left-lateralized network of regions relevant for language processing and learning, as well as ancillary processes such as attentional control, including lIFG, IAnIns, rAnIns, lSMA, lSPL, lTPJ, and lFuG. These regions have been consistently implicated in lexical and grammatical learning, phonological processing, integration of novel word meanings, and attentional control during vocabulary acquisition via explicit or implicit learning mechanisms (Breitenstein & Knecht 2002; Breitenstein et al. 2005; Rodriguez-Fornells et al., 2009; Laine & Salmelin, 2009; Tagarelli et al., 2019).

Notably, the learning task was adapted based on a previous study (Sliwinska et al., 2017) and despite several design modifications (e.g., increased number of learning blocks, visual vs. auditory pseudoword presentation, continuous vs. sparse fMRI acquisition), both studies showed similar outcomes. Behaviorally, the time course and degree of improvement across learning stages were comparable, underscoring the robustness of APPL as a word-learning paradigm. Importantly, task-evoked activity and changes across learning stages largely overlapped, with more pronounced task-evoked activity in early learning stages and a gradual decrease when proficiency increased. This convergence supports the view that the identified regions are most strongly engaged during early, effortful encoding, with reduced recruitment once associations are consolidated. Such decreases in neural activity with repeated exposure are consistent with findings from pseudoword repetition studies (e.g., Rauschecker et al., 2008), which demonstrate learning-related reductions in activation in left inferior frontal, superior temporal, and motor-related regions. These effects are commonly interpreted as increased neural efficiency or sharpening of stimulus representations, reflecting more efficient perceptual–motor and phonological processing of newly learned word forms.

For example, it has been suggested that the MDC and domain-specific regions interact to provide a temporary neural module that enables rapid learning and outcome monitoring (Engström et al. 2013; Ruge & Wolfensteller 2016). With increased task proficiency, the role of the MDC is thought to decrease, likely reflecting decreased demands on central cognitive resources (e.g., working memory, selective attention, or performance monitoring) that support early stages of associative learning (Chein & Schneider 2012). The specificity of this pattern to early stages of learning is also in line with results for the lexical decision control task that elicited less pronounced activity in MDC (i.e., preSMA, bilaterally in the insula) and showed no gradual decrease over time. Notably, learning has also been linked to decreased activity in domain specific brain regions (e.g., Breitenstein et al. 2005; Vartanian et al. 2022), possibly related to sharpening of representations and changes of within- and between network activity. Decreased activity in bilateral IFG has been linked to more effortful processing or increased selection demands (Meinzer et al., 2009; Meinzer et al., 2012), and improved word-retrieval during IFG-tDCS was associated with decreased activity in bilateral IFG (Meinzer et al., 2012; 2013). While correlational, linking this overall pattern of decreased activity to behavioral learning success further extends previous findings by Slivinska et al. (2017), and results highlight the specificity and functional relevance of effects observed during APPL. It is acknowledged that the exact neurophysiological mechanisms underlying short-term learning (and potential tDCS effects) and their relation to fMRI derived metrics are not well understood and cannot be derived from this study. For further information, we refer the readers to recent in-depth reviews of current theoretical accounts and tentative mechanisms (e.g., Fertonani & Miniussi 2017; Gazerani 2025).

The present study also addresses contemporary methodological debates about how to inform valid and reliable stimulation targets for noninvasive brain stimulation in cognitive neuroscience (see Meinzer et al. 2024). For novel-word learning, previous tDCS studies have mainly stimulated core language regions in a uniform (i.e., non-individualized) fashion, based on theoretical models of language learning or imaging meta-analyses involving similar tasks (e.g., Flöel et al., 2008; Meinzer et al., 2014; Perceval et al., 2017; Tagarelli et al., 2019; Kurmakaeva et al., 2021; Perikova et al., 2022). However, empirical evidence for the effectiveness of this approach remained mixed, and several studies have reported relatively weak or inconsistent tDCS effects (for a recent discussion, see Riemann et al., 2024), possibly related to suboptimal target regions for tDCS. Importantly, due to the neuromodulatory nature of tDCS, its effectiveness critically depends on the overlap of the induced current with brain activity elicited by specific paradigms and in specific populations (Meinzer et al., 2024). Moreover, effects in brain stimulation studies can be obscured by limited TRR of experimental paradigms (Falleti et al., 2006; Hausknecht et al., 2007; Bartels et al., 2010) or imaging biomarkers (Elliott et al., 2020; Compère et al., 2021), i.e., confounds that have rarely been considered in previous tDCS research (Abdelmotaleb et al., 2025).

This was directly addressed in the present study by identifying valid, functionally relevant, and reliable target regions for a subsequent crossover tDCS-fMRI study. Specifically, data analyses confirmed the stability of behavioral outcomes that showed good-to-excellent consistency across sessions. Functional imaging TRR analyses demonstrated variability of ICCs even within task-active regions, with a median ICC of 0.696. However, a voxel-wise analysis of our activation patterns also revealed that ∼81% of voxels achieved moderate-to-excellent reliability (see **Supplementary Figure 4**). This was complemented by two ROI analyses demonstrating that the majority of a priori ROIs (including left IFG, SMA, and SPL) and all peak ICC ROIs showed good-to-excellent consistency. Power to detect true effects with our sample size was also confirmed for the majority of ROIs. Overall, these results compare favorably to a recent meta-analysis of cognitive fMRI studies, which demonstrated relatively poor TRR across repeated sessions, with an average ICC of 0.397 and substantial variability within ROIs (Elliott et al., 2020). Nonetheless, the observed results in our study are also in line with suggestions that reliability estimates tend to diminish when moving beyond localized peaks to broader activation patterns. Hence, the identified cortical activity peaks with high TRR in our study are interpreted as core processing nodes during the APPL task, rendering them as primary targets for neurostimulation targets. These included lIFG, ISMA, and ISPL as particularly reliable and behaviorally relevant, while another region that was frequently targeted in language-learning studies (lTPJ; e.g., Flöel 2008; Perceval et al. 2017) showed good TRR but weaker activity and correlation with learning. Future studies involving active tDCS arms are required to investigate which of the identified candidate regions are optimal for enhancing novel word learning.

Our findings furthermore converge with and extend those of Abdelmotaleb et al. (2025), who used the same overall design structure to investigate potential target regions for enhancing object–location memory (OLM). This study reported good to excellent reliability of behavioral outcomes and the highest TRR for peak activity in key nodes of the OLM network. While this study did not report voxelwise TRR analyses, visual inspection of their data suggests a similar pattern as in our own. Taken together, the results indicate that both paradigms are well-suited for crossover tDCS–fMRI studies, providing a robust framework for detecting stimulation effects in future studies that employ the same tasks with demographically matched participants. However, our study also outlines a general path towards optimizing brain stimulation studies by implementing an empirically informed approach to selecting valid and functionally relevant target regions and by implementing reliable behavioral and imaging paradigms.

## 5. Limitations and Future Directions

Despite promising results, some limitations need to be acknowledged: Behaviorally, we observed ceiling effects for accuracy in later stages that may reduce sensitivity to detect improvements from tDCS. The latter suggests that RT will be the superior choice for future active tDCS studies using the APPL paradigm. Moreover, there was a numerically small, but significant main effect of task version. Hence, counterbalancing of stimulus sets is advised, which is especially important in cross-over tDCS studies to minimize potential biases related to stimulus properties. The relatively small sample size limits the precision of TRR estimates. Power analyses suggested valid results for the majority of the identified target regions, but this requires confirmation in larger and independent samples. Moreover, imaging outcomes are study specific and may not generalize to other populations with structural and functional network reorganization (e.g., healthy older individuals, patients). We also demonstrated in our post-hoc modeling analyses that a simple projection of cortical target coordinates to the scalp for montage selection can result in meaningful current intensities in the underlying cortex. However, readers are cautioned that the reported modeling outcomes are also study specific and future studies need to consider further optimizations (for discussion see Meinzer et al. 2024; Thielscher et al. 2026), including prospective, individualized current modeling and neuronavigated electrode placement (Niemann et al., 2024). The latter ensures that electrodes are attached as intended, which is particularly relevant for focal tDCS setups that deliver the current to relative circumscribed brain regions. In addition, we relied on group level coordinates for targeting and individual peaks may vary across participants. In practice, this limitation is likely mitigated by the fact that even focal tDCS still induces neurophysiologically relevant current flow to a relatively large area (see **Figure 7**). Finally, using fMRI to identify functionally relevant target regions is time and cost-intensive and may not always be feasible. However, testing the stability of behavioral measures in pre-tDCS experiments is recommended as a minimal approach.

## 6. Conclusions

In sum, our findings demonstrate that the adapted APPL task is a valid and reliable paradigm for investigating novel-word learning in future tDCS-fMRI studies. Identification of valid, reliable, and functionally relevant target regions has potential for enhancing the effectiveness of tDCS in the future, and the suggested approach can serve as a blueprint for optimizing the use of this method to enhance other cognitive functions.

## Funding

The author(s) declare that financial support was received for the research, authorship, and/or publication of this article. This research was funded by the German Research Foundation (project grants: CRC INST 276/741-2 and 292/155-1, Research Unit 5429/1 (467143400), ME 3161/5-1, ME 3161/6-1, AN1103/5-1; FL 379/34-1); Heisenberg grant to DA AN1103/6-1 [539593253].

## Ethics Statement

The study was directed at the University Medicine Greifswald and permitted by the medical ethics committee (Registry number BB015/22) and this study followed the guidelines of the Declaration of Helsinki, and subjects provided printed informed consent beforehand inclusion.

## Acknowledgments

This study was performed in the context of the multi-center study MeMoSLAP - “Modulation of brain networks for memory and learning by transcranial electrical brain stimulation: A systematic, lifespan approach”; Research Unit: 5429, funded by the German Research Foundation (Deutsche Forschungsgemeinschaft, DFG).

## Declaration of interests

The authors declare that they have no known competing financial interests or personal relationships that could have appeared to influence the work reported in this paper.

## Data Availability Statement

The study was pre-registered with the Open Science Framework, and the protocol can be accessed through OSF Registries (https://osf.io/t37u2). The corresponding author can provide the data supporting the study’s conclusions upon request. Due to ethical and privacy concerns about the study’s participants, the data are not publicly accessible.

## Notes

### Competing Interest Statement

The authors have declared no competing interest.

### Summary of Updates

Following the major revision requested by NeuroImage, we conducted a few additional analyses and updated existing analyses accordingly. As a result, most figures have been revised, Table 1 has been removed, and new supplementary methods, figures and tables have been added. In addition: 1. We revised the Methods section to improve clarity and consistency regarding the ROI analyses described in Sections 3.4.2 and 3.5. 2. We corrected an issue in the task based analyses and updated the corresponding results. The main findings and brain and behavior relationships remained consistent with the original submission, while the test retest reliability results showed improvement.

## References

Abdelmotaleb, M., Niemann, F., Kocataş, H., Caisachana Guevara, L. M., Shahbabaie, A., Malinowski, R., Riemann, S., Fromm, A. E., Hayek, D., Antonenko, D., Meinzer, M., & Flöel, A. (2025). Identification of Reliable Target Brain Regions for Enhancing Object–Location Memory by Brain Stimulation. Brain and Behavior, 15(7). 10.1002/brb3.70658

Antal, A., Bjekić, J., Ganho-Ávila, A., Alekseichuk, I., Assecondi, S., Bergmann, T. O., Bikson, M., Brunelin, J., Brunoni, A. R., Charvet, L., Chen, R., Cohen Kadosh, R., Diedrich, L., D’Urso, G., Ferrucci, R., Filipović, S. R., Fitzgerald, P. B., Flöel, A., Fröhlich, F., … Ziemann, U. (2026). Low intensity transcranial electric stimulation: Safety, ethical, legal regulatory and application guidelines (2017-2025: An update) - endorsed by the European Society for Brain Stimulation (ESBS) and by the International Federation for Clinical Neurophysiology (IFCN). Clinical Neurophysiology: Official Journal of the International Federation of Clinical Neurophysiology, 184, 2111436. 10.1016/j.clinph.2025.2111436

Bakdash, J. Z., & Marusich, L. R. (2017). Repeated Measures Correlation. Frontiers in Psychology, 8, 456. 10.3389/fpsyg.2017.00456

Bartels, C., Wegrzyn, M., Wiedl, A., Ackermann, V., & Ehrenreich, H. (2010). Practice effects in healthy adults: A longitudinal study on frequent repetitive cognitive testing. BMC Neuroscience, 11(1), 118. 10.1186/1471-2202-11-118

Breitenstein, C., & Knecht, S. (2002). Development and validation of a language learning model for behavioral and functional-imaging studies. Journal of Neuroscience Methods, 114(2), 173–179. 10.1016/S0165-0270(01)00525-8

Breitenstein, C., Jansen, A., Deppe, M., Foerster, A.-F., Sommer, J., Wolbers, T., & Knecht, S. (2005). Hippocampus activity differentiates good from poor learners of a novel lexicon. NeuroImage, 25(3), 958–968. 10.1016/j.neuroimage.2004.12.019

Caceres, A., Hall, D. L., Zelaya, F. O., Williams, S. C. R., & Mehta, M. A. (2009). Measuring fMRI reliability with the intra-class correlation coefficient. NeuroImage, 45(3), 758–768. 10.1016/j.neuroimage.2008.12.035

Chein, J. M., & Schneider, W. (2012). The Brain’s Learning and Control Architecture. Current Directions in Psychological Science, 21(2), 78–84. 10.1177/0963721411434977

Compère, L., Siegle, G. J., & Young, K. (2021). Importance of test–retest reliability for promoting fMRI based screening and interventions in major depressive disorder. Translational Psychiatry, 11(1), 387. 10.1038/s41398-021-01507-3

Demidenko, M., Mumford, J., & Russ Poldrack. (2024). PyReliMRI: An Open-source Python tool for Estimates of Reliability in MRI Data (Version 2.1.0) [Software]. Zenodo. 10.5281/ZENODO.12522260

Donner, A., & Eliasziw, M. (1987). Sample size requirements for reliability studies. Statistics in Medicine, 6(4), 441–448. 10.1002/sim.4780060404

Duñabeitia, J. A., Baciero, A., Antoniou, K., Antoniou, M., Ataman, E., Baus, C., Ben-Shachar, M., Çağlar, O. C., Chromý, J., Comesaña, M., Filip, M., Đurđević, D. F., Dowens, M. G., Hatzidaki, A., Januška, J., Jusoh, Z., Kanj, R., Kim, S. Y., Kırkıcı, B., … Pliatsikas, C. (2022). The Multilingual Picture Database. Scientific Data, 9(1), 431. 10.1038/s41597-022-01552-7

Elliott, M. L., Knodt, A. R., Ireland, D., Morris, M. L., Poulton, R., Ramrakha, S., Sison, M. L., Moffitt, T. E., Caspi, A., & Hariri, A. R. (2020). What Is the Test-Retest Reliability of Common Task-Functional MRI Measures? New Empirical Evidence and a Meta-Analysis. Psychological Science, 31(7), 792–806. 10.1177/0956797620916786

Engstrom, M., Landtblom, A.-M., & Karlsson, T. (2013). Brain and effort: Brain activation and effort-related working memory in healthy participants and patients with working memory deficits. Frontiers in Human Neuroscience, 7. 10.3389/fnhum.2013.00140

Falleti, M. G., Maruff, P., Collie, A., & Darby, D. G. (2006). Practice Effects Associated with the Repeated Assessment of Cognitive Function Using the CogState Battery at 10-minute, One Week and One Month Test-retest Intervals. Journal of Clinical and Experimental Neuropsychology, 28(7), 1095–1112. 10.1080/13803390500205718

Fertonani, A., & Miniussi, C. (n.d.). Transcranial Electrical Stimulation: What We Know and Do Not Know About Mechanisms.

Flöel, A., Rösser, N., Michka, O., Knecht, S., & Breitenstein, C. (2008). Noninvasive brain stimulation improves language learning. Journal of Cognitive Neuroscience, 20(8), 1415–1422. 10.1162/jocn.2008.20098

Gazerani, P. (2025). The neuroplastic brain: Current breakthroughs and emerging frontiers. Brain Research, 1858, 149643. 10.1016/j.brainres.2025.149643

Grady, C. L., Rieck, J. R., Nichol, D., Rodrigue, K. M., & Kennedy, K. M. (2020). Influence of sample size and analytic approach on stability and interpretation of brain-behavior correlations in task-related fMRI data. Human Brain Mapping, 42(1), 204–219. 10.1002/hbm.25217

Greenberg, E., & Macias, R. F. (o. J.). English Literacy and Language Minorities in the United States. Education Statistics Quarterly, 3(4), 73e75.

Guo, J., Zhang, K., Zhang, J., Zhao, R., Liang, Y., Lin, Y., Yu, S., Qin, W., & Yang, X. (2021). Decoding Spatial Memory Retrieval in Cubical Space Using fMRI Signals. Frontiers in Neural Circuits, 15, 624352. 10.3389/fncir.2021.624352

Harth, S., & Müller, S. V. (2006). Testrezension. Zeitschrift für Neuropsychologie. https://econtent.hogrefe.com/doi/10.1024/1016-264X.15.4.315

Hausknecht, J. P., Halpert, J. A., Di Paolo, N. T., & Moriarty Gerrard, M. O. (2007). Retesting in selection: A meta-analysis of coaching and practice effects for tests of cognitive ability. Journal of Applied Psychology, 92(2), 373–385. 10.1037/0021-9010.92.2.373

Honda, M., Deiber, M. P., Ibáñez, V., Pascual-Leone, A., Zhuang, P., & Hallett, M. (1998). Dynamic cortical involvement in implicit and explicit motor sequence learning. A PET study. Brain: A Journal of Neurology, 121 ( *Pt 11*), 2159–2173. 10.1093/brain/121.11.2159

Kaestle, C. F., Campbell, A., Finn, J. D., Johnson, S. T., & Mikulecky, L. J. (2001). Adult Literacy and Education in America: Four Studies Based on the National Adult Literacy Survey. ED Pubs, P. https://eric.ed.gov/?id=ED461718

Köhler, S., Moscovitch, M., Winocur, G., Houle, S., & McIntosh, A. R. (1998). Networks of domain-specific and general regions involved in episodic memory for spatial location and object identity. Neuropsychologia, 36(2), 129–142. 10.1016/s0028-3932(97)00098-5

Koo, T. K., & Li, M. Y. (2016). A Guideline of Selecting and Reporting Intraclass Correlation Coefficients for Reliability Research. Journal of Chiropractic Medicine, 15(2), 155–163. 10.1016/j.jcm.2016.02.012

Kühner, C., Bürger, C., Keller, F., & Hautzinger, M. (2007). Reliabilität und Validität des revidierten Beck-Depressionsinventars (BDI-II): Befunde aus deutschsprachigen Stichproben. Der Nervenarzt, 78(6), 651–656. 10.1007/s00115-006-2098-7

Kurmakaeva, D., Blagovechtchenski, E., Gnedykh, D., Mkrtychian, N., Kostromina, S., & Shtyrov, Y. (2021). Acquisition of concrete and abstract words is modulated by tDCS of Wernicke’s area. Scientific Reports, 11(1), 1508. 10.1038/s41598-020-79967-8

Laine, M., & Salmelin, R. (2010). Neurocognition of New Word Learning in the Native Tongue: Lessons From the Ancient Farming Equipment Paradigm. Language Learning, 60(s2), 25–44. 10.1111/j.1467-9922.2010.00599.x

Lehrl, S. (2018). Multiple-Choice Vocabulary Test, MWT. (6. Auflage). Spitta GmbH.

Lo, S., & Andrews, S. (2015). To transform or not to transform: Using generalized linear mixed models to analyse reaction time data. Frontiers in Psychology, 6. 10.3389/fpsyg.2015.01171

McGraw, K. O., & Wong, S. P. (1996). Forming inferences about some intraclass correlation coefficients. Psychological Methods, 1(1), 30–46. 10.1037/1082-989X.1.1.30

Meinzer, M., Antonenko, D., Lindenberg, R., Hetzer, S., Ulm, L., Avirame, K., Flaisch, T., & Flöel, A. (2012). Electrical Brain Stimulation Improves Cognitive Performance by Modulating Functional Connectivity and Task-Specific Activation. Journal of Neuroscience, 32(5), 1859–1866. 10.1523/JNEUROSCI.4812-11.2012

Meinzer, M., Flaisch, T., Seeds, L., Harnish, S., Antonenko, D., Witte, V., Lindenberg, R., & Crosson, B. (2012). Same Modulation but Different Starting Points: Performance Modulates Age Differences in Inferior Frontal Cortex Activity during Word-Retrieval. PLOS ONE, 7(3), e33631. 10.1371/journal.pone.0033631

Meinzer, M., Flaisch, T., Wilser, L., Eulitz, C., Rockstroh, B., Conway, T., Rothi, L., & Crosson, B. (2009). Neural Signatures of Semantic and Phonemic Fluency in Young and Old Adults. Journal of Cognitive Neuroscience, 21, 2007–2018. 10.1162/jocn.2009.21219

Meinzer, M., Jähnigen, S., Copland, D. A., Darkow, R., Grittner, U., Avirame, K., Rodriguez, A. D., Lindenberg, R., & Flöel, A. (2014). Transcranial direct current stimulation over multiple days improves learning and maintenance of a novel vocabulary. Cortex, 50, 137–147. 10.1016/j.cortex.2013.07.013

Meinzer, M., Lindenberg, R., Antonenko, D., Flaisch, T., & Flöel, A. (2013). Anodal transcranial direct current stimulation temporarily reverses age-associated cognitive decline and functional brain activity changes. The Journal of Neuroscience: The Official Journal of the Society for Neuroscience, 33(30), 12470–12478. 10.1523/JNEUROSCI.5743-12.2013

Meinzer, M., Lindenberg, R., Darkow, R., Ulm, L., Copland, D., & Flöel, A. (2014). Transcranial Direct Current Stimulation and Simultaneous Functional Magnetic Resonance Imaging. Journal of Visualized Experiments, 86, 51730. 10.3791/51730

Meinzer, M., Shahbabaie, A., Antonenko, D., Blankenburg, F., Fischer, R., Hartwigsen, G., Nitsche, M. A., Li, S.-C., Thielscher, A., Timmann, D., Waltemath, D., Abdelmotaleb, M., Kocataş, H., Caisachana Guevara, L. M., Batsikadze, G., Grundei, M., Cunha, T., Hayek, D., Turker, S., … Flöel, A. (2024). Investigating the neural mechanisms of transcranial direct current stimulation effects on human cognition: Current issues and potential solutions. Frontiers in Neuroscience, 18, 1389651. 10.3389/fnins.2024.1389651

Müller, H., Hasse-Sander, I., Horn, R., Helmstaedter, C., & Elger, C. E. (1997). Rey auditory–verbal learning test: Structure of a modified German version. Journal of Clinical Psychology, 53(7), 663–671. 10.1002/(SICI)1097-4679(199711)53:7%253C663::AID-JCLP4%253E3.0.CO;2-J

Niemann, F., Shababaie, A., Paßmann, S., Riemann, S., Malinowski, R., Kocataş, H., Caisachana Guevara, L. M., Abdelmotaleb, M., Antonenko, D., Blankenburg, F., Fischer, R., Hartwigsen, G., Li, S.-C., Nitsche, M. A., Thielscher, A., Timmann, D., Fromm, A., Hayek, D., Hubert, A.-K., … Meinzer, M. (2024). Neuronavigated Focalized Transcranial Direct Current Stimulation Administered During Functional Magnetic Resonance Imaging. Journal of Visualized Experiments, 213, 67155. 10.3791/67155

Noble, S., Scheinost, D., & Constable, R. T. (2021). A guide to the measurement and interpretation of fMRI test-retest reliability. Current Opinion in Behavioral Sciences, 40, 27–32. 10.1016/j.cobeha.2020.12.012

Oldfield, R. C. (1971). The assessment and analysis of handedness: The Edinburgh inventory. Neuropsychologia, 9(1), 97–113. 10.1016/0028-3932(71)90067-4

Perceval, G., Martin, A. K., Copland, D. A., Laine, M., & Meinzer, M. (2017). High-definition tDCS of the temporo-parietal cortex enhances access to newly learned words. Scientific Reports, 7(1), 17023. 10.1038/s41598-017-17279-0

Perceval, G., Martin, A. K., Copland, D. A., Laine, M., & Meinzer, M. (2020). Multisession transcranial direct current stimulation facilitates verbal learning and memory consolidation in young and older adults. Brain and Language, 205, 104788. 10.1016/j.bandl.2020.104788

Perikova, E., Blagovechtchenski, E., Filippova, M., Shcherbakova, O., Kirsanov, A., & Shtyrov, Y. (2022). Anodal tDCS over Broca’s area improves fast mapping and explicit encoding of novel vocabulary. Neuropsychologia, 168, 108156. 10.1016/j.neuropsychologia.2022.108156

Petersson, K. M., Elfgren, C., & Ingvar, M. (1999). Dynamic changes in the functional anatomy of the human brain during recall of abstract designs related to practice. Neuropsychologia, 37(5), 567–587. 10.1016/s0028-3932(98)00152-3

Poldrack, R. A. (2007). Region of interest analysis for fMRI. Social Cognitive and Affective Neuroscience, 2(1), 67–70. 10.1093/scan/nsm006

Ramos-Escobar, N., Segura, E., Olive, G., Rodriguez-Fornells, A., & François, C. (2021). Oscillatory activity and EEG phase synchrony of concurrent word segmentation and meaning-mapping in 9-year-old children. Developmental Cognitive Neuroscience, 51, 101010. 10.1016/j.dcn.2021.101010

Rauschecker, A. M., Pringle, A., & Watkins, K. E. (2008). Changes in neural activity associated with learning to articulate novel auditory pseudowords by covert repetition. Human Brain Mapping, 29(11), 1231–1242. 10.1002/hbm.20460

Riemann, S., Van Lück, J., Rodríguez-Fornells, A., Flöel, A., & Meinzer, M. (2024). The role of frontal cortex in novel-word learning and consolidation: Evidence from focal transcranial direct current stimulation. Cortex, 177, 15–27. 10.1016/j.cortex.2024.05.004

Ripollés, P., Biel, D., Peñaloza, C., Kaufmann, J., Marco-Pallarés, J., Noesselt, T., & Rodríguez-Fornells, A. (2017). Strength of Temporal White Matter Pathways Predicts Semantic Learning. The Journal of Neuroscience, 37(46), 11101–11113. 10.1523/JNEUROSCI.1720-17.2017

Rodríguez-Fornells, A., Cunillera, T., Mestres-Missé, A., & De Diego-Balaguer, R. (2009). Neurophysiological mechanisms involved in language learning in adults. Philosophical Transactions of the Royal Society B: Biological Sciences, 364(1536), 3711–3735. 10.1098/rstb.2009.0130

Ruge, H., & Wolfensteller, U. (2016). Towards an understanding of the neural dynamics of intentional learning: Considering the timescale. NeuroImage, 142, 668–673. 10.1016/j.neuroimage.2016.06.006

Setsompop, K., Gagoski, B. A., Polimeni, J. R., Witzel, T., Wedeen, V. J., & Wald, L. L. (2012). Blipped-controlled aliasing in parallel imaging for simultaneous multislice echo planar imaging with reduced *g* -factor penalty. Magnetic Resonance in Medicine, 67(5), 1210–1224. 10.1002/mrm.23097

Shtyrov, Y. (2012). Neural Bases of Rapid Word Learning. The Neuroscientist, 18(4), 312–319. 10.1177/1073858411420299

Sliwinska, M. W., Violante, I. R., Wise, R. J. S., Leech, R., Devlin, J. T., Geranmayeh, F., & Hampshire, A. (2017). Stimulating Multiple-Demand Cortex Enhances Vocabulary Learning. The Journal of Neuroscience, 37(32), 7606–7618. 10.1523/JNEUROSCI.3857-16.2017

Tagarelli, K. M., Shattuck, K. F., Turkeltaub, P. E., & Ullman, M. T. (2019). Language learning in the adult brain: A neuroanatomical meta-analysis of lexical and grammatical learning. NeuroImage, 193, 178–200. 10.1016/j.neuroimage.2019.02.061

Thielscher, A., Antunes, A., & Saturnino, G. B. (2015). Field modeling for transcranial magnetic stimulation: A useful tool to understand the physiological effects of TMS? Annual International Conference of the IEEE Engineering in Medicine and Biology Society. IEEE Engineering in Medicine and Biology Society. Annual International Conference, 2015, 222–225. 10.1109/EMBC.2015.7318340

Thielscher, A., Hayek, D., Puonti, O., Grittner, U., Blankenburg, F., Fischer, R., Hartwigsen, G., Li, S.-C., Meinzer, M., Nitsche, M. A., Timmann, D., Flöel, A., & Antonenko, D. (2026). Harmonizing the stimulation dose of focal transcranial direct current stimulation across target sites. NeuroImage, 331, 121882. 10.1016/j.neuroimage.2026.121882

Tombaugh, T. (2004). Trail Making Test A and B: Normative data stratified by age and education. Archives of Clinical Neuropsychology, 19(2), 203–214. 10.1016/S0887-6177(03)00039-8

Tong, Y., Chen, Q., Nichols, T. E., Rasetti, R., Callicott, J. H., Berman, K. F., Weinberger, D. R., & Mattay, V. S. (2016). Seeking Optimal Region-Of-Interest (ROI) Single-Value Summary Measures for fMRI Studies in Imaging Genetics. PLOS ONE, 11(3), e0151391. 10.1371/journal.pone.0151391

Turner, B. O., Paul, E. J., Miller, M. B., & Barbey, A. K. (2018). Small sample sizes reduce the replicability of task-based fMRI studies. Communications Biology, 1(1), 62. 10.1038/s42003-018-0073-z

Van Der Elst, W., Van Boxtel, M. P. J., Van Breukelen, G. J. P., & Jolles, J. (2006). The Stroop Color-Word Test: Influence of Age, Sex, and Education; and Normative Data for a Large Sample Across the Adult Age Range. Assessment, 13(1), 62–79. 10.1177/1073191105283427

Vartanian, O., Replete, V., Saint, S. A., Lam, Q., Forbes, S., Beaudoin, M. E., Brunyé, T. T., Bryant, D. J., Feltman, K. A., Heaton, K. J., McKinley, R. A., Van Erp, J. B. F., Vergin, A., & Whittaker, A. (2022). What Is Targeted When We Train Working Memory? Evidence From a Meta-Analysis of the Neural Correlates of Working Memory Training Using Activation Likelihood Estimation. Frontiers in Psychology, 13. 10.3389/fpsyg.2022.868001

Walter, S. D., Eliasziw, M., & Donner, A. (1998). Sample size and optimal designs for reliability studies. Statistics in Medicine, 17(1), 101–110. 10.1002/(sici)1097-0258(19980115)17:1%3C101::aid-sim727%3E3.0.co;2-e

Weir, J. P. (2005). Quantifying Test-Retest Reliability Using the Intraclass Correlation Coefficient and the SEM. The Journal of Strength and Conditioning Research, 19(1), 231. 10.1519/15184.1

Woods, D. L., Kishiyama, M. M., Yund, E. W., Herron, T. J., Edwards, B., Poliva, O., Hink, R. F., & Reed, B. (2011). Improving digit span assessment of short-term verbal memory. Journal of Clinical and Experimental Neuropsychology, 33(1), 101–111. 10.1080/13803395.2010.493149

Xu, J., Moeller, S., Auerbach, E. J., Strupp, J., Smith, S. M., Feinberg, D. A., Yacoub, E., & Uğurbil, K. (2013). Evaluation of slice accelerations using multiband echo planar imaging at 3 T. NeuroImage, 83, 991–1001. 10.1016/j.neuroimage.2013.07.055

Yarkoni, T. (2009). Big Correlations in Little Studies: Inflated fMRI Correlations Reflect Low Statistical Power-Commentary on Vul et al. (2009). Perspectives on Psychological Science: A Journal of the Association for Psychological Science, 4(3), 294–298. 10.1111/j.1745-6924.2009.01127.x

Young, A. R., Beitchman, J. H., Johnson, C., Douglas, L., Atkinson, L., Escobar, M., & Wilson, B. (2002). Young adult academic outcomes in a longitudinal sample of early identified language impaired and control children. Journal of Child Psychology and Psychiatry, 43(5), 635–645. 10.1111/1469-7610.00052

Yucel, A., Niemann, F., Meinzer, M., & Martin, A. K. (2025). Regionally specific picture naming benefits of focal tDCS are dependent on baseline performance in older adults. GeroScience. 10.1007/s11357-025-01674-x

Zhang, X., Lv, L., Min, G., Wang, Q., Zhao, Y., & Li, Y. (2021). Overview of the Complex Figure Test and Its Clinical Application in Neuropsychiatric Disorders, Including Copying and Recall. Frontiers in Neurology, 12. 10.3389/fneur.2021.680474

