## Supplementary_Materials_R1 for "Functionally Relevant and Reliable Brain Stimulation Targets for Enhancement of Novel Word-Learning"

### Supplementary Figures

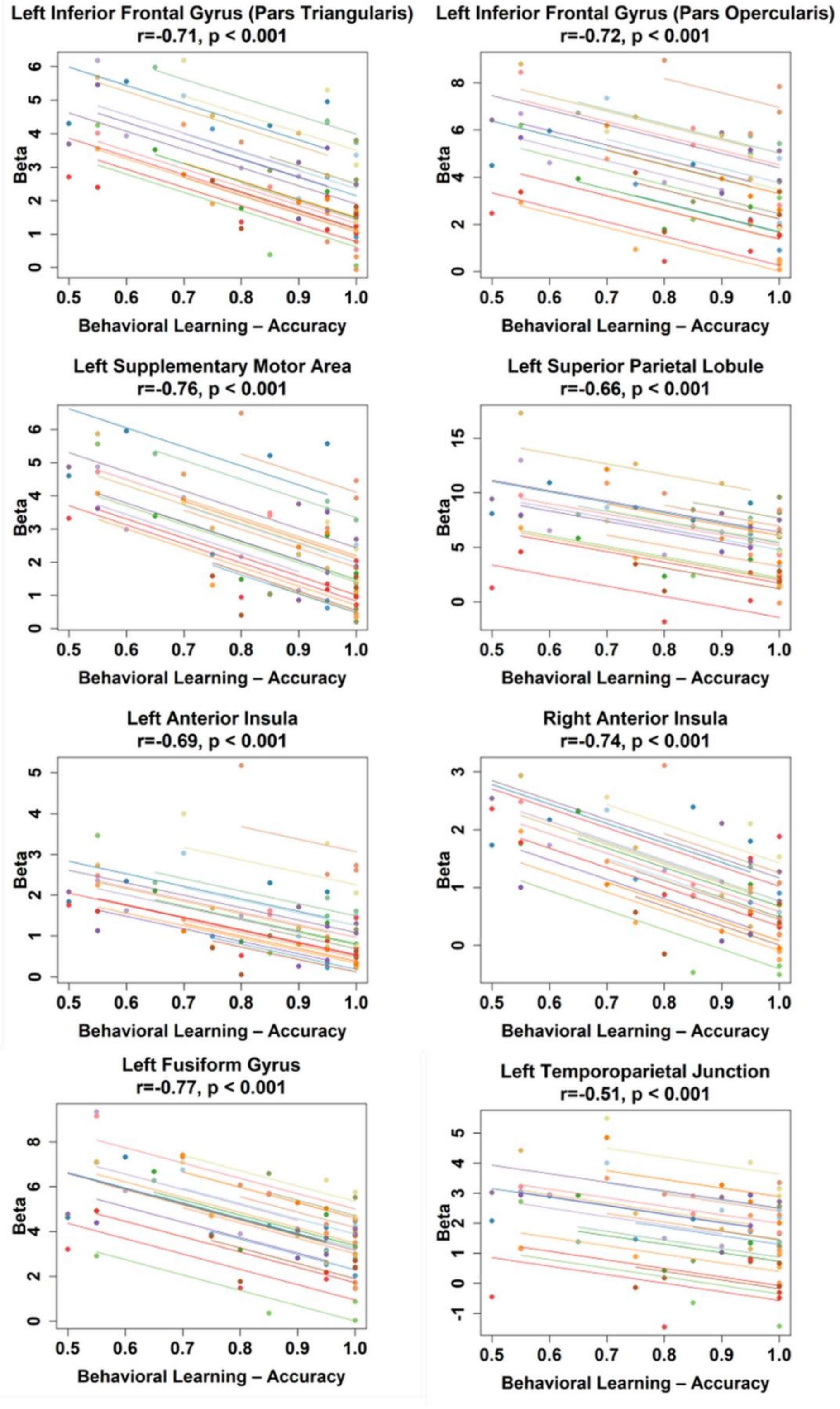


**Supplementary Figure 1: Repeated-measures correlations between behavioral learning performance (accuracy) and brain activity (beta values) across learning stages in a priori regions-of-interest (ROIs).** Points represent participants and stages (20 participants, 4 stages), pooled across the two imaging sessions. Lines represent individual participant-specific trends across the four stages (n = 20), illustrating within-subject trajectories. Increased accuracy across learning stages was associated with lower beta values. Correlation coefficients (r) and associated p-values are reported for each ROI. lIFGpt: Left Inferior Frontal Gyrus (Pars Triangularis), lIFGpo: Left Inferior Frontal Gyrus (Pars Opercularis), lSMA: Left Supplementary Motor Area, lSPL: Left Superior Parietal Lobule, lAnIns: Left Anterior Insula, rAnIns: Right Anterior Insula, lFuG: Left Fusiform Gyrus, lTPJ: Left Temporoparietal Junction.

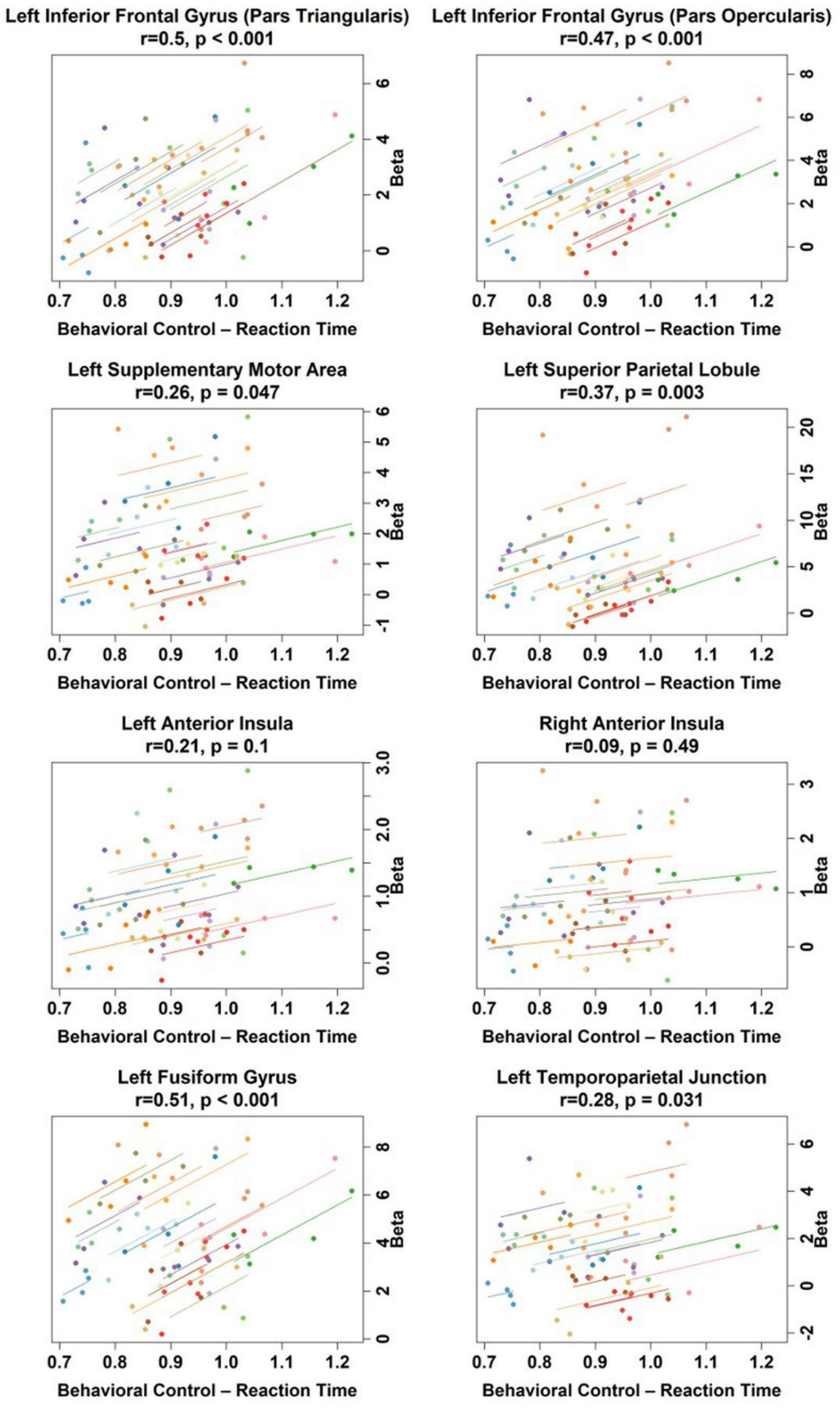


**Supplementary Figure 2: Repeated-measures correlations between behavioral control task performance (reaction time, RT) and brain activity (beta values) across stages in a priori regions-of-interest (ROIs).** Points represent participants and stages (20 participants, 4 stages), pooled across the two imaging sessions. Lines represent individual participant-specific trends across the four stages (n = 20), illustrating within-subject trajectories. Shorter RT (faster response latency) was associated with lower beta values. Correlation coefficients (r) and associated p-values are reported for each ROI. Note: The y-axis is plotted on the right to emphasize the decrease in RT across the four stages lIFGpt: Left Inferior Frontal Gyrus (Pars Triangularis), lIFGpo: Left Inferior Frontal Gyrus (Pars Opercularis), lSMA: Left Supplementary Motor Area, lSPL: Left Superior Parietal Lobule, lAnIns: Left Anterior Insula, rAnIns: Right Anterior Insula, lFuG: Left Fusiform Gyrus, lTPJ: Left Temporoparietal Junction.

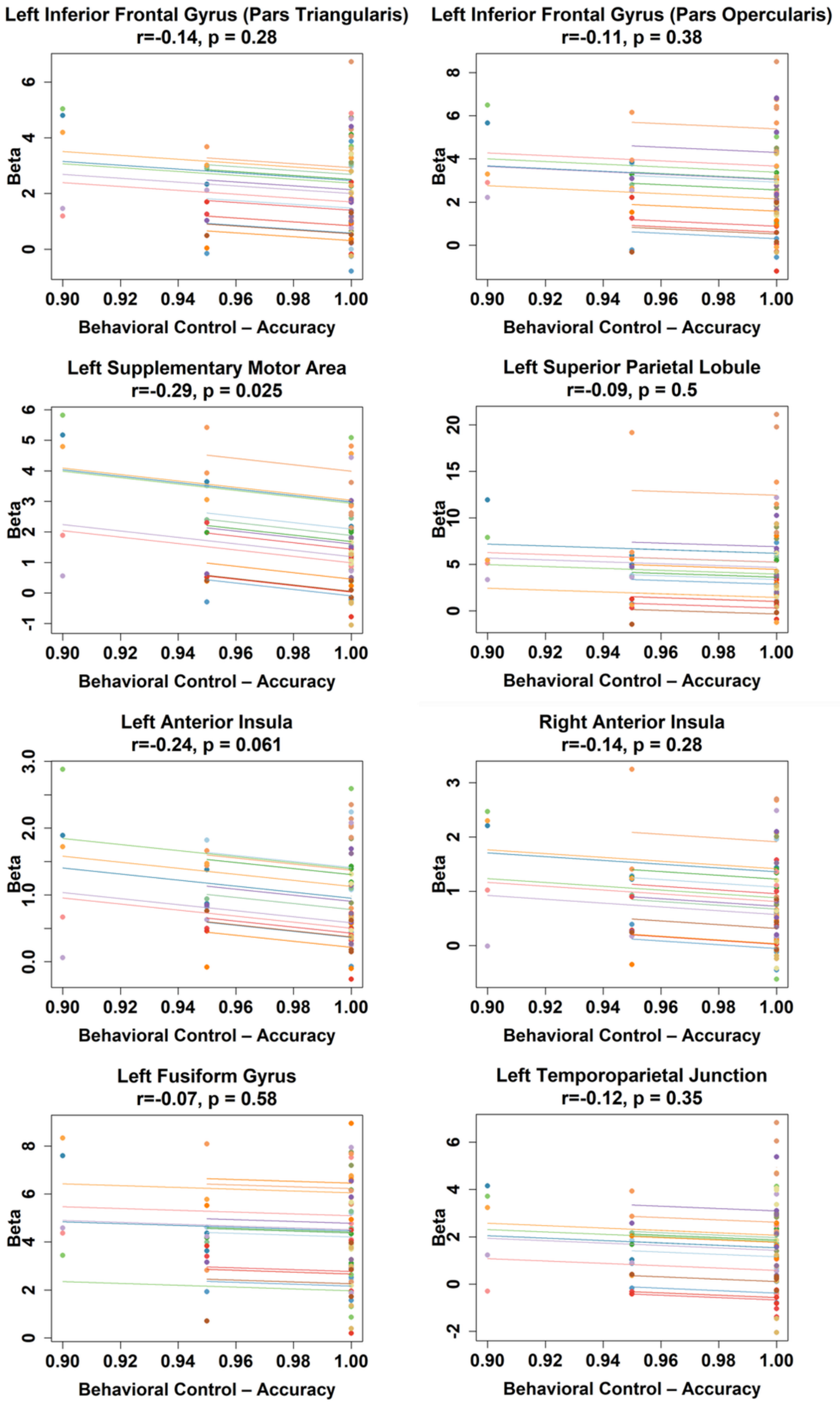


**Supplementary Figure 3: Repeated-measures correlations between behavioral control task performance (accuracy) and brain activity (beta values) across learning stages in a priori regions-of-interest (ROIs).** Points represent participants and stages (20 participants, 4 stages), pooled across the two imaging sessions. Lines represent individual participant-specific trends across the four stages (n = 20), illustrating within-subject trajectories. Correlation coefficients (r) and associated p-values are reported for each ROI. lIFGpt: Left Inferior Frontal Gyrus (Pars Triangularis), lIFGpo: Left Inferior Frontal Gyrus (Pars Opercularis), lSMA: Left Supplementary Motor Area, lSPL: Left Superior Parietal Lobule, lAnIns: Left Anterior Insula, rAnIns: Right Anterior Insula, lFuG: Left Fusiform Gyrus, lTPJ: Left Temporoparietal Junction.

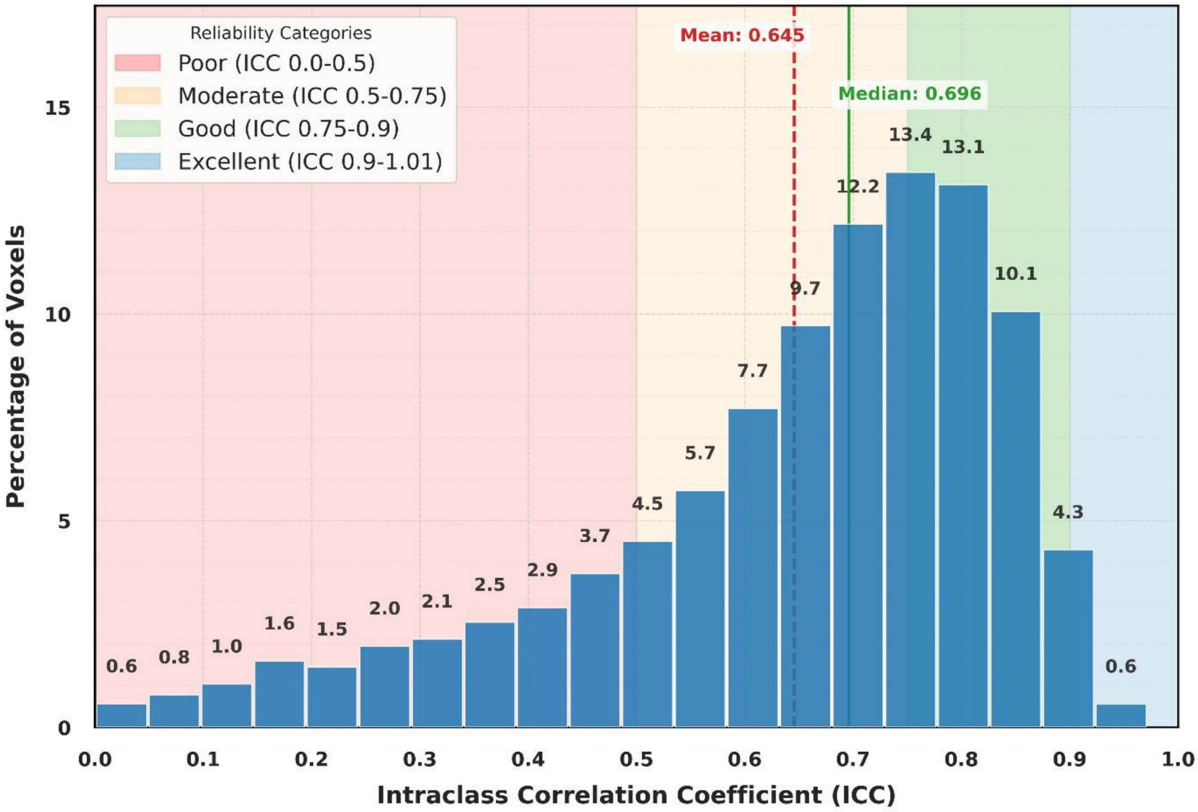


**Supplementary Figure 4: Test-retest reliability within the task-related network.** Histogram of voxel-wise ICC values in the APPL-related activation (N = 28,866 voxels). Bars show the percentage of voxels per ICC bin (20 intervals), with percentages labeled above. Background colors indicate reliability ranges as described in the legend. APPL= associative picture-pseudoword learning. ICC= Intraclass Correlation Coefficient.

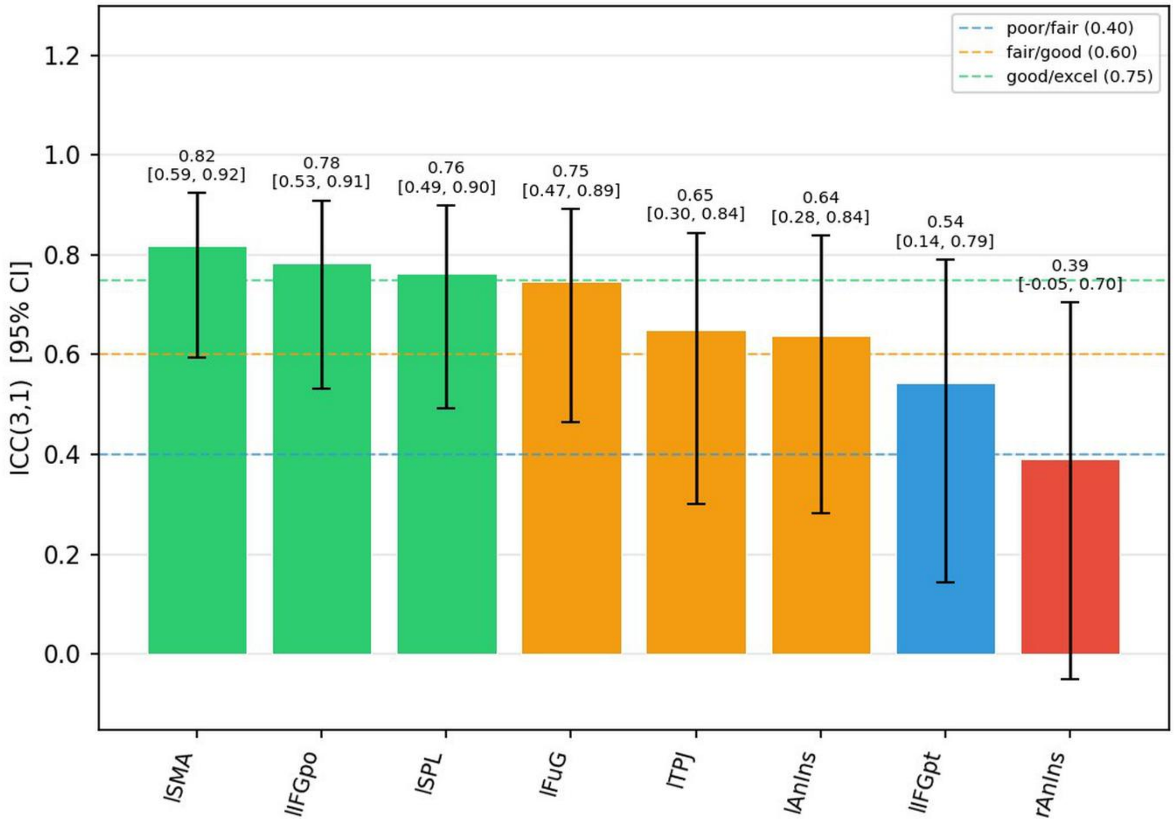


**Supplementary Figure 5**: **ICC Summary Bar Chart (A Priori ROIs)**. ICC (3,1) estimates with 95% confidence intervals for the eight a priori ROIs. Bar colors indicate ICC quality thresholds: green (≥ 0.75, good/excellent), orange (≥ 0.60, fair/good), blue (≥ 0.40, poor/fair), red (< 0.40, poor). Dashed horizontal lines mark the quality thresholds. Numerical annotations above each bar show the point estimate and 95% CI.

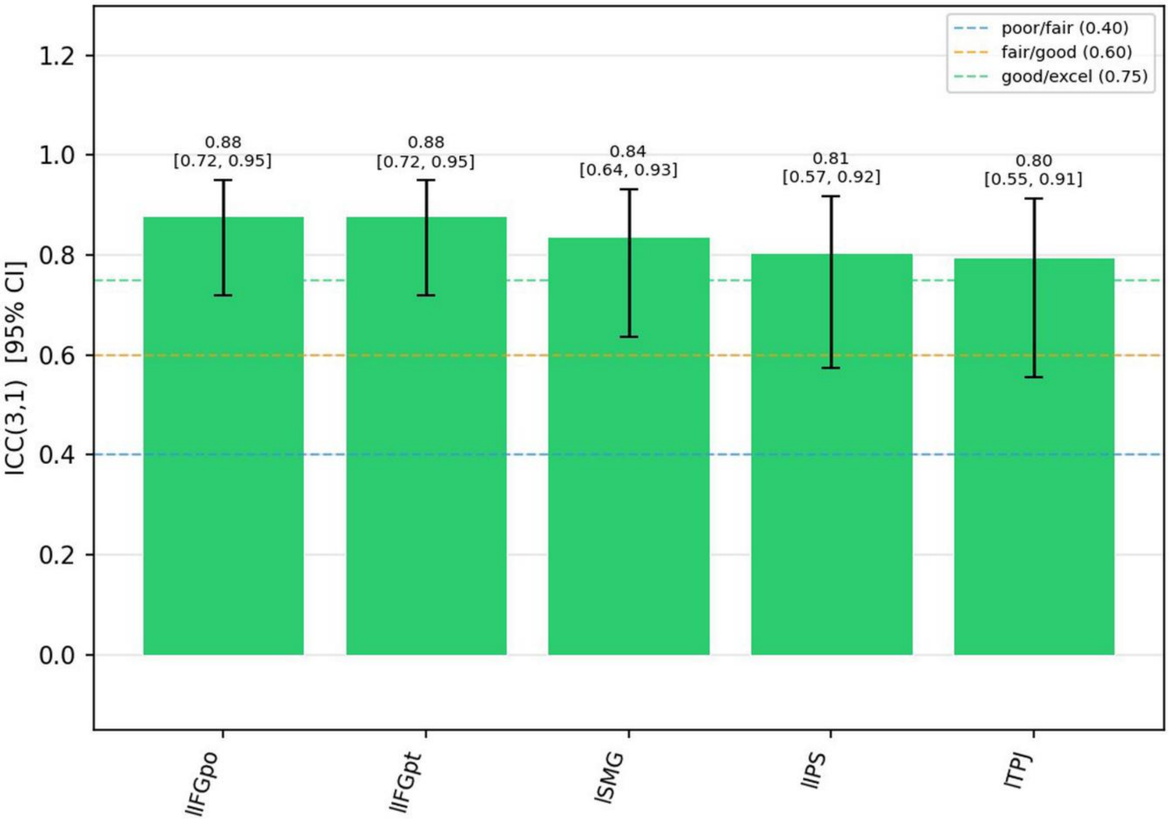


**Supplementary Figure 6**: **ICC Summary Bar Chart (ICC-Peak ROIs)**. ICC (3,1) estimates with 95% confidence intervals for the five ICC-peak ROIs. Conventions as above. All ROIs exceeded the good/excellent threshold (ICC ≥ 0.75).

### Supplementary Tables

| Variable | Mean: Set 1 | SD: Set 1 | Mean: Set 2 | SD: Set 2 | p-value |
|---|---|---|---|---|---|
| **APPL Task** | | | | | |
| Pseudoword length (letters) | 5.64 | 0.93 | 5.50 | 0.87 | 0.5625 |
| Object name length (letters) | 5.43 | 0.94 | 5.64 | 1.26 | 0.4821 |
| Word frequency (Simple Celex) | 91.00 | 137.19 | 92.36 | 123.36 | 0.9657 |
| Modal name usage (%) | 92.54 | 13.25 | 92.84 | 10.58 | 0.9274 |
| Visual complexity | 2.52 | 0.53 | 2.58 | 0.60 | 0.6965 |
| **Control Task** | | | | | |
| Word Length (letters) | 6.45 | 1.50 | 5.75 | 0.94 | 0.0930 |
| Word Frequency (Simple Celex) | 89.00 | 88.62 | 87.75 | 57.79 | 0.9445 |
| Modal Name Usage (%) | 91.22 | 10.87 | 91.18 | 12.24 | 0.9918 |
| Visual Complexity | 2.64 | 0.67 | 2.42 | 0.44 | 0.2349 |

**Supplementary Table 1:** Matching parameters and statistical comparison of stimulus sets used for the associative picture-pseudoword learning (APPL) and control tasks.

**Fixed effects from the accuracy model (Binomial GLMM)**

| Parameter | β | SE | z value | p-value | 95% CI |
|---|---|---|---|---|---|
| **Intercept** | 3.781 | 0.289 | 13.10 | <.001 | [3.215, 4.347] |
| **Stage** | -0.072 | 0.089 | -0.80 | .421 | [-0.247, 0.103] |
| **Condition (learning)** | -4.049 | 0.262 | -15.44 | <.001 | [-4.563, -3.535] |
| **Session 2** | -0.044 | 0.063 | -0.70 | .486 | [-0.168, 0.080] |
| **Task version (2)** | 0.208 | 0.063 | 3.29 | .001 | [0.084, 0.332] |
| **Block** | -0.030 | 0.013 | -2.27 | .023 | [-0.056, -0.004] |
| **Stage × Condition (learning)** | 1.210 | 0.098 | 12.40 | <.001 | [1.018, 1.402] |

**Fixed effects from the reaction time model (Inverse Gaussian GLMM)**

| Parameter | β | SE | t value | p-value | 95% CI |
|---|---|---|---|---|---|
| **Intercept** | 2.3083 | 0.0324 | 71.18 | <.001 | [2.244, 2.372] |
| **Stage** | -0.0210 | 0.0030 | -7.07 | <.001 | [-0.027, -0.015] |
| **Condition (learning)** | 0.2202 | 0.0100 | 22.09 | <.001 | [0.200, 0.240] |
| **Session 2** | -0.0382 | 0.0036 | -10.70 | <.001 | [-0.045, -0.031] |
| **Task version (2)** | -0.0010 | 0.0036 | -0.28 | .784 | [-0.008, 0.006] |
| **Block** | -0.0073 | 0.0008 | -9.43 | <.001 | [-0.0089, -0.0058] |
| **Stage × Condition (learning)** | -0.0572 | 0.0035 | -16.15 | <.001 | [-0.064, -0.050] |

**Supplementary Table 2: Fixed effects from accuracy and reaction time models;** these tables summarize the fixed effects from the accuracy (Binomial GLMM) and reaction time (Inverse Gaussian GLMM) models. For each effect, the estimate (β), standard error (SE), z- or t-value, and p-value are reported. The models included all stages, conditions, sessions, task types, and relevant interactions.

| Brain Region | Side | Cluster Size (voxels) | T-value | X | Y | Z |
|---|---|---|---|---|---|---|
| Supramarginal Gyrus, posterior division | L | 1017 | 8.74 | -44 | -42 | 48 |
| Lateral Occipital Cortex, superior division | L | | 6.60 | -24 | -64 | 32 |
| Superior Parietal Lobule | L | | 6.16 | -32 | -54 | 48 |
| Juxtapositional Lobule Cortex (Supplementary Motor Cortex) | L | 1127 | 7.89 | -4 | 0 | 62 |
| Paracingulate Gyrus | L | | 6.21 | -2 | 14 | 48 |
| Paracingulate Gyrus | R | | 6.14 | 8 | 10 | 50 |
| Precentral Gyrus | L | 1054 | 7.77 | -50 | -8 | 42 |
| | | | 7.19 | -40 | -4 | 48 |
| | | | 6.29 | -50 | 0 | 48 |
| Insula Cortex | L | 299 | 6.48 | -30 | 22 | -2 |
| Temporal Fusiform Cortex, posterior division | R | 192 | 6.11 | 32 | 22 | 10 |
| Superior Frontal Gyrus | R | | 4.94 | 36 | 18 | -2 |

**Supplementary Table 3: Harvard-Oxford cortical-subcortical structural atlas.** ***Whole brain comparison of the APPL > control tasks.*** Significance threshold $p < 0.001$ (uncorrected), with a cluster-based family-wise error correction (FWEc). Cluster extent >192. Hemisphere (Right = R & Left = L), cluster size in voxels, peak coordinates in MNI space, X, Y, Z). Brain regions were labeled using the Harvard–Oxford cortical and subcortical structural atlas.

**Supplementary Table 4: Power Analysis (A Priori ROIs)**

| ROI | ICC (3,1) | Achieved Power | N for 80% Power |
|---|---|---|---|
| lSMA | 0.817 | 1.000 | 5 |
| lIFGpo | 0.785 | 1.000 | 6 |
| lSPL | 0.763 | 1.000 | 6 |
| lFuG | 0.748 | 1.000 | 7 |
| lTPJ | 0.649 | 0.989 | 10 |
| lAnIns | 0.638 | 0.984 | 10 |
| lIFGpt | 0.543 | 0.898 | 16 |
| rAnIns | 0.390 | 0.586 | 35 |

**Note:** "Achieved Power" = power to reject $H_0$: ICC = 0 at $\alpha = 0.05$ with N = 20 and k = 2 sessions. "N for 80% Power" = minimum sample size required to achieve ≥ 80% power.

**Supplementary Table 5: Power Analysis (ICC-Peak ROIs)**

| ROI | ICC (3,1) | Achieved Power | N for 80% Power |
|---|---|---|---|
| lIFGpo | 0.879 | 1.000 | 5 |
| lIFGpt | 0.879 | 1.000 | 5 |
| lSMG | 0.838 | 1.000 | 5 |
| lIPS | 0.806 | 1.000 | 5 |
| lTPJ | 0.796 | 1.000 | 5 |

**Note:** "Achieved Power" = power to reject $H_0$: ICC = 0 at $\alpha = 0.05$ with N = 20 and k = 2 sessions. "N for 80% Power" = minimum sample size required to achieve ≥ 80% power.

### Supplementary Methods

#### Electric Field Modeling for Target Translation

A critical challenge for translating fMRI-identified cortical targets into practical stimulation protocols (i.e., tDCS set-ups) requires an understanding of how electrode arrangements and placement over the target regions relates to the induced electric field distribution.

This supplementary analysis is aimed to bridge the gap between mapping target regions and designing practical tDCS montages by modeling electric fields for representative target regions. For illustrative purposes, we chose three functionally relevant cortical target regions identified in the main manuscript and calculated their closest scalp projection point (for details see Niemann et al., 2024). This served as starting point for implementing 3x1 montages (as described in the main manuscript) and electrical field simulations aimed at determining the intensity and spatial extent of the induced current (1) across the whole brain and (2) in regions-of-interest centered around the target regions. Numerous other approaches are possible (e.g., see Thielscher et al., 2026) and researchers are advised to implement modeling approaches best suited for their specific study requirements. Notably, individualized field simulations as described in the main manuscript are study specific and the outcomes may not directly translate into optimal montages for other studies or populations.

#### Methods

Individual head models were constructed for our 20 study participants using the nonlinear warp from charm (mni2subject_coords, SimNIBS 4.5, Thielscher et al., 2015). T1- and T2-weighted MRI scans of each participant were segmented into five tissue compartments (scalp, skull, CSF, grey matter, white matter) using the charm pipeline with default conductivities. 3×1 focal tDCS montages were modeled, targeting the selected functionally relevant left-hemispheric regions (i.e., corresponding to the a priori ROIs described in the main manuscript: lIFG triangularis: MNI [−46, 28, 20]; lTPJ: MNI [−60, −43, 22]; lSMA: MNI [−4, 6, 60]).

Details of the individualized current modeling procedure are described in Niemann et al. (2024). In brief, for each montage, target MNI coordinates were transformed to individual head space using the nonlinear warp from charm (mni2subject_coords). A circular anode (20 mm diameter, 2 mm thickness) was placed on the scalp by transforming the target MNI coordinate to the individual subject’s scalp surface using the nonlinear warp from the charm pipeline. To define positions of the three surround cathodes, we utilized a customized SimNIBS Python function (get_surround_pos and expand_to_center_surround), which generated the focal tDCS configuration based on the individual's anatomy. This function extracted the local scalp geometry around the anode and calculated a best-fit sphere to the skin ROI to establish a local coordinate system. The three cathodes were then projected onto the individual scalp mesh at 120° intervals along a 40 mm center-to-center radius. This geometry-aware approach ensured that the 3×1 configuration maintained precise center-to-center distances while conforming to the unique cranial curvature of each participant. A total current of 2 mA (anode: +2 mA; each cathode: −2/3 mA) was modeled. Finite element method simulations computed electric field magnitude (|E|, V/m) for each subject and montage. Results were mapped to FsAverage cortical surface for group visualization. To quantify field strength in each target, we also defined 10 mm radius spherical ROIs centered on the transformed target coordinate in each subject's gray matter volume. Bcause the tetrahedric mesh volumes have a different size for each volumes, volume-weighted mean |E|were computed for each ROI.

#### ROI-Specific ICC Analyses and Power Analysis

Intraclass correlation coefficient (ICC) analyses were calculated for (a) a priori ROIs derived from a recent neuroanatomical meta-analysis of lexical learning (Tagarelli et al., 2019) and (b) ICC-peak ROIs identified from the voxel-wise ICC map of the present study. These analyses complement the whole-brain ICC maps by quantifying reliability at the ROI level and providing formal power analyses to assess the adequacy of the current sample size (N = 20) for detecting reliable ICC effects.

All ICC estimates were computed using ICC (3,1), a two-way mixed-effects model with absolute consistency for single measurements (McGraw & Wong, 1996), implemented via the PyReliMRI toolbox (Demidenko et al., 2024). Mean beta values were extracted within 5 mm radius spheres centered on the specified MNI coordinates. Power analyses follow the non-central F approximation for ICC reliability studies (Donner & Eliasziw, 1987; Walter et al., 1998).

Spherical ROIs (5 mm radius) were centered on MNI coordinates reported in the meta-analysis. ICC (3,1) was computed from session-wise mean beta values (learning > implicit baseline) extracted for each participant in the two fMRI sessions. Peak ICC coordinates are reported in the main manuscript.

#### ICC-Peak ROI Analysis

To complement the a priori approach, we also examined reliability at peak-voxel coordinates identified from the voxel-wise ICC map of the APPL task (see Figure 6 of the main manuscript). These coordinates represent voxels with the highest local ICC values within clusters of task-related activation. Spherical ROIs (5 mm radius) were centered on these ICC-peak coordinates and subjected to the same analysis pipeline described for the a priori ROIs.

#### Power Analysis

To address concerns about sample size adequacy, we performed a formal power analysis for each ROI. Power was estimated using the non-central F approximation for the ICC F-test under a two-way mixed model with $k=2$ sessions and $\alpha=0.05$ (Donner & Eliasziw, 1987; Walter et al., 1998). For each ROI, we report the achieved power with the current sample size (N=20).

#### References


Demidenko, M., Mumford, J., & Russ Poldrack. (2024). *PyReliMRI: An Open-source Python tool for Estimates of Reliability in MRI Data* (Version 2.1.0) [Computer software]. Zenodo. https://doi.org/10.5281/ZENODO.12522260

Donner, A., & Eliasziw, M. (1987). Sample size requirements for reliability studies. *Statistics in Medicine, 6*(4), 441–448. https://doi.org/10.1002/sim.4780060404

McGraw, K. O., & Wong, S. P. (1996). Forming inferences about some intraclass correlation coefficients. *Psychological Methods, 1*(1), 30–46. https://doi.org/10.1037/1082-989X.1.1.30

Niemann, F., Shababaie, A., Paßmann, S., Riemann, S., Malinowski, R., Kocataş, H., Caisachana Guevara, L. M., Abdelmotaleb, M., Antonenko, D., Blankenburg, F., Fischer, R., Hartwigsen, G., Li, S.-C., Nitsche, M. A., Thielscher, A., Timmann, D., Fromm, A., Hayek, D., Hubert, A.-K., … Meinzer, M. (2024). Neuronavigated Focalized Transcranial Direct Current

Stimulation Administered During Functional Magnetic Resonance Imaging. *Journal of Visualized Experiments*, (213), 67155. https://doi.org/10.3791/67155

Tagarelli, K. M., Shattuck, K. F., Turkeltaub, P. E., & Ullman, M. T. (2019). Language learning in the adult brain: A neuroanatomical meta-analysis of lexical and grammatical learning. *NeuroImage*, *193*, 178–200. https://doi.org/10.1016/j.neuroimage.2019.02.061

Thielscher, A., Antunes, A., & Saturnino, G. B. (2015). Field modeling for transcranial magnetic stimulation: A useful tool to understand the physiological effects of TMS? *Annual International Conference of the IEEE Engineering in Medicine and Biology Society. IEEE Engineering in Medicine and Biology Society. Annual International Conference*, *2015*, 222–225. https://doi.org/10.1109/EMBC.2015.7318340

Thielscher, A., Hayek, D., Puonti, O., Grittner, U., Blankenburg, F., Fischer, R., Hartwigsen, G., Li, S.-C., Meinzer, M., Nitsche, M. A., Timmann, D., Flöel, A., & Antonenko, D. (2026). Harmonizing the stimulation dose of focal transcranial direct current stimulation across target sites. *NeuroImage*, *331*, 121882. https://doi.org/10.1016/j.neuroimage.2026.121882

Walter, S. D., Eliasziw, M., & Donner, A. (1998). Sample size and optimal designs for reliability studies. *Statistics in Medicine*, *17*(1), 101–110. https://doi.org/10.1002/(sici)1097-0258(19980115)17:1%3C101::aid-sim727%3E3.0.co;2-e
